# Climate change negatively impacts dominant microbes in the sediments of a High Arctic lake

**DOI:** 10.1101/705178

**Authors:** Graham A. Colby, Matti O. Ruuskanen, Kyra A. St. Pierre, Vincent L. St. Louis, Alexandre J. Poulain, Stéphane Aris-Brosou

## Abstract

Temperatures in the Arctic are expected to increase dramatically over the next century, yet little is known about how microbial communities and their underlying metabolic processes will be affected by these environmental changes in freshwater sedimentary systems. To address this knowledge gap, we analyzed sediments from Lake Hazen, NU Canada. Here, we exploit the spatial heterogeneity created by varying runoff regimes across the watershed of this uniquely large lake at these latitudes to test how a transition from low to high runoff, used as one proxy for climate change, affects the community structure and functional potential of dominant microbes. Based on metagenomic analyses of lake sediments along these spatial gradients, we show that increasing runoff leads to a decrease in taxonomic and functional diversity of sediment microbes. Our findings are likely to apply to other, smaller, glacierized watersheds typical of polar or high latitude / high altitudes ecosystems; we can predict that such changes will have far reaching consequences on these ecosystems by affecting nutrient biogeochemical cycling, the direction and magnitude of which are yet to be determined.

## Introduction

Climate change is amplified in polar regions, where near-surface temperatures have increased almost twice as fast as elsewhere on Earth over the last decade (Overpeck *et al.*, 1997; Serreze and Francis, 2006; Screen and Simmonds, 2010). Climate models predict that temperature will increase in the Arctic by as much as 8°C by 2100 (IPCC, 2013). These changes are already having dramatic consequences on physical (Laudon *et al.*, 2017; Bliss *et al.*, 2014; O’Reilly *et al.*, 2015), biogeochemical (Frey and McClelland, 2009; Lehnherr *et al.*, 2018), and ecological (Smol *et al.*, 2005; Wrona *et al.*, 2016) processes across Arctic ecosystems. Yet, while we are starting to understand the effect of thawing permafrost on microbial communities (McCalley *et al.*, 2014; Hultman *et al.*, 2015; Mackelprang *et al.*, 2016), we know very little about how microbes in lentic ecosystems such as lakes respond to environmental changes – even though microbes mediate most global biogeochemical cycles (Falkowski *et al.*, 2008; Fuhrman, 2009). Furthermore, lakes are broadly considered sentinels of climate change, as they integrate physical, chemical and biological changes happening through their watersheds (Williamson *et al.*, 2009); however, their microbial community structure and function are relatively understudied, in particular in the Arctic.

To date, much of the research performed on microbial communities in Arctic lakes has been limited to studies that were mostly based on partial 16S rRNA gene sequencing (Stoeva *et al.*, 2014; Thaler *et al.*, 2017; Mohit *et al.*, 2017; Ruuskanen *et al.*, 2018a; Cavaco *et al.*, 2019). While these studies are useful to understand the structure of these microbial communities, they provide limited functional insights and can be biased as they often rely on sequence databases where environmental microbes, specifically from the Arctic, may be underrepresented (Ruuskanen *et al.*, 2018b, 2019). More critically, being circumscribed both in space and in time, previous studies only offer snapshots of microbial communities and hence, have a limited power to predict how microbial communities might respond to climate change.

To predict the effect of climate change on microbial functional diversity in Arctic lake sediments, we focused on Lake Hazen, the world’s largest High Arctic lake by volume (82°N) (Köck *et al.*, 2012). In this work, we exploited two important properties of Lake Hazen. First, its watershed is already experiencing the effects of climate change, as increasing temperatures there are leading to more glacial melt, permafrost thaw, and increased runoff from the watershed into the lake in warmer years relative to cooler ones (Lehnherr *et al.*, 2018). Second, its tributaries are highly heterogeneous, fed by eleven glaciers ranging from 6 to 1041 km^2^ in surface area, and annual runoff volume approximately scaling with their size (from *<*0.001 to 0.080 km^3^ in 2016) (Pierre *et al.*, 2019).

It is this temporal and spatial heterogeneity in runoff that we used to evaluate the possible consequences of climate change on High Arctic sediment microbial functional diversity, acknowledging that the consequences of increasing temperature are likely slightly more plural and complex. To this effect, we sampled lake sediments along two transects representing low (L transect: samples L1 [shallow] and L2 [deep]) and high (H: samples H1 [shallow] and H2, [deep]) seasonal runoff volume, as well as at a single site that received negligible runoff (C site; Figure 1A). We also collected soil samples (S sites) from three sites in the dried streambeds of the tributaries, on the northern shore between the two transects to assess soil influence on microbial communities present in the sediments. We then resorted to untargeted metagenomics analyses to draw an inventory of dominant microbes, assumed to be the most critical to nutrient cycling and the most relevant to the dynamics of microbial communities. These reconstructed Metagenome Assembled Genomes (MAGs) (Bowers *et al.*, 2017) allowed us to assess the quantitative impact of a change of runoff regime, from low to high, on both the structure of sediment microbial communities and their functional potential. We show that an increase in runoff volume and resultant sedimentation rates, as predicted under climate change scenarios for the region, could lead to a reduced diversity of the dominant microbial community and of their functional potential.

**Figure 1.**
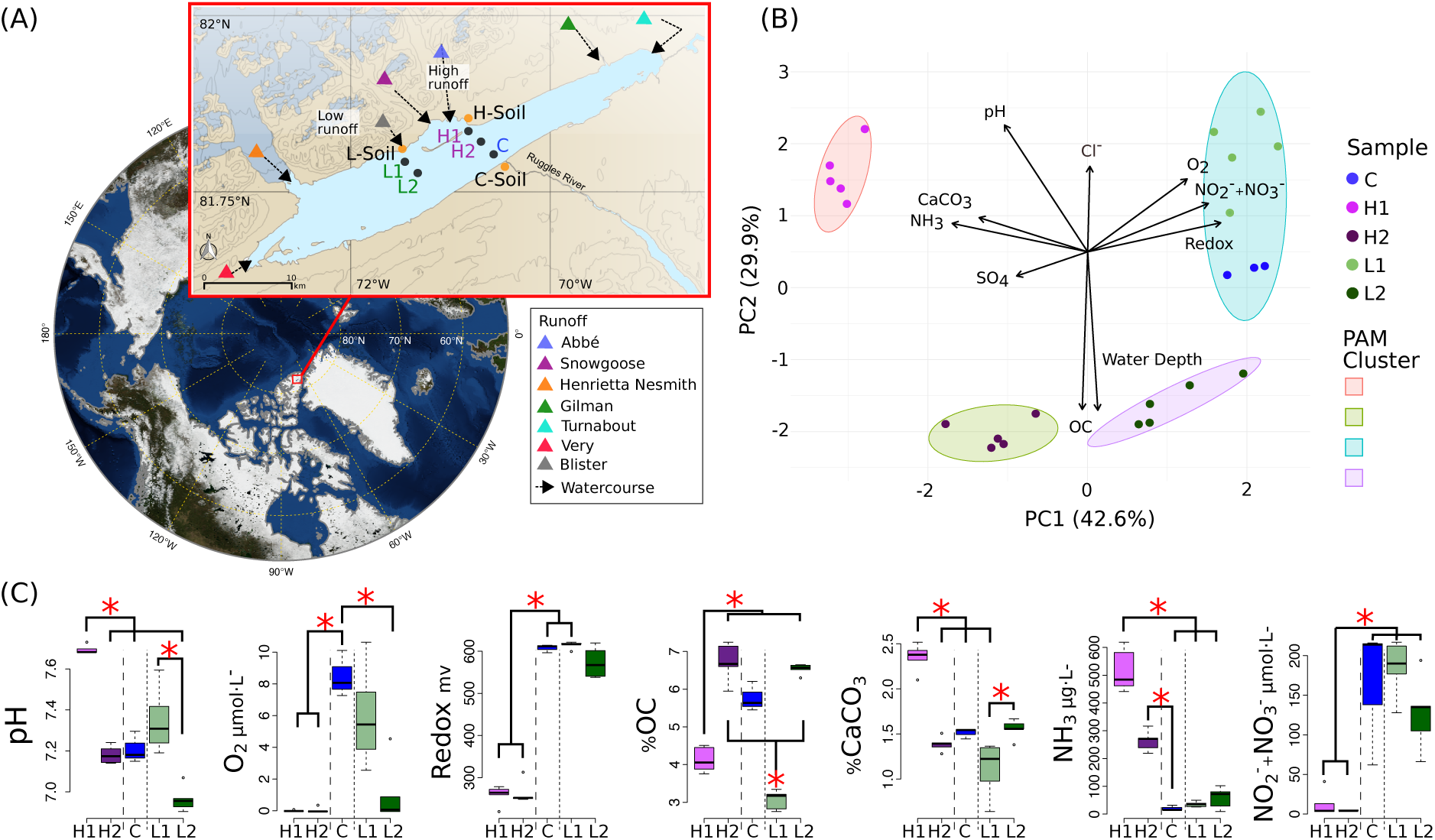
Lake Hazen sampling design and chemical composition. (A) Location of Lake Hazen (red box). Inset map: soil (orange dots) and sediment (black dots) sample sites are separated into hydrological regimes of high (purple), low (green), and negligible/control (blue) runoff. (B) Principal component analysis (PCA) showing the differences in physical and chemical composition of the sediment sites. Vectors display pH, dissolved dioxygen (O_2_), redox potential, nitrates and nitrites concentration 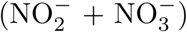, water depth, percent organic carbon (OC), percent calcium carbonate (CaCO_3_), sulfate 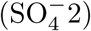 concentration (SO_4_), and ammonia concentration (NH_3_). Individual points represent the mean values using 1 cm intervals measured in the top 5 cm. Partitioning around medoids was used to identify clusters. (C) Distribution of chemical features for sediment sites. Branches and asterisks indicate significant differences between sites *P <* 0.025 (Dunn Test). If branch tips form a dichotomy or trichotomy, the interactions within that group is not significant. Long dashes separate high runoff sites and dotted line separates low runoff sites. There was insufficient data to include soil sites in B and C.

## Methods

### Sample collection and processing

Sediment and soil cores were collected from Lake Hazen (82°N, 71°W: Figure 1A), located within Quttinirpaaq National Park, on northern Ellesmere Island, Nunavut. Sampling took place between May 10 and June 10, 2017, when the lake was still completely ice-covered (Supplementary Table 1). Within the watershed, runoff flows from the outlet glaciers along the northwestern shoreline through poorly consolidated river valleys, depositing sediments at the bottom of Lake Hazen along two transects, the H1/H2 and L1/L2 sites, respectively. The lake then drains via the Ruggles River along its south-eastern shoreline (C sites). The surrounding glacial rivers deliver different amounts of sediments, nutrients and organic carbon unevenly to the lake as a consequence of heterogeneous sedimentation rates (Supplementary Table 2). More specifically, the top 5 cm of sediments from the deeper low (L2) and high (H2) runoff sites represented depositional periods of 30 years (1987-2017) and 6 years (2011-2017), respectively (Supplementary Table 3).

Samples were collected along two transects and can be separated into three hydrological regimes by seasonal runoff volume: low (L transect), high (H transect), and negligible runoff (C sites) summarized in Supplementary Table 3. Contamination of samples was minimized by wearing non-powdered latex gloves during sample handling and sterilizing all equipment with 10% bleach and 90% ethanol before sample collection. Sediment cores approximately 30 cm in length were collected with an UWITEC (Mondsee, Austria) gravity corer from five locations: C (overlying water depth: 50 m) far from the direct influence of glacial inflows serving as a control site; L1 (water depth: 50 m) and L2 (water depth: 251 m), at variable distances from a small glacial inflow (Blister Creek, *<*0.001 km^3^ in summer 2016); and, H1 (water depth: 21 m) and H2 (water depth: 253 m), located adjacent to several larger glacial inflows (*i.e.*, the Abbé River, 0.015 km^3^ and Snow Goose, 0.006 km^3^ in 2016). The soil samples (S sites) were collected from three sites in the dried streambeds of the tributaries, on the northern shore between the two transects. At each site, for both sediments and soil, five cores were sampled, ∼3 m apart for the sediment cores, and approximately ∼1 m apart to account for site heterogeneity.

For sediment core, one of the five cores were used for microprofiling of oxygen (O_2_), redox and pH, as well as one core for porewater chemistry and loss on ignition (see (Ruuskanen *et al.*, 2018a) for details), and the remaining three cores were combined, prior to their genomic analysis, here again to account for site heterogeneity. For soil samples, three cores per site were collected for DNA analysis, but no additional cores were collected for chemical analyses. As we were mostly interested in the community composition through space, we combined the top 5 cm of sediment and 10 cm of soil for sample extraction and subsequent sequencing. Any remaining length of cores that were used for DNA analysis were discarded. These uppermost layers in the sediment correspond to both the most recent sediment deposition dates (Pierre *et al.*, 2019) and the region of greatest microbial activity (Haglund *et al.*, 2003). The top of each core was sectioned and placed into Whirlpack bags. These slices were homogenized manually inside of the bags and stored in a −20°C freezer until shipment back to the University of Ottawa where they were then stored at −80°C. Soil samples were transferred into falcon tubes, homogenized, and stored as described above for the lake sediment samples.

Samples were thawed overnight and 250-400 mg (wet weight; Supplementary Table 4) were then washed in a sterile salt buffer (10 mM EDTA, 50 mM Tris-HCl, 50 mM Na_2_ HPO_4_ 7H_2_O at pH 8.0) to remove PCR inhibitors (Zhou *et al.*, 1996; Poulain *et al.*, 2015). All sample handling was conducted in a stainless-steel laminar flow hood (HEPA 100) treated with UVC radiation and bleach before use. DNA extractions were performed using the DNeasy PowerSoil Kit (MO BIO Laboratories Inc, Carlsbad, CA, USA), following the kit guidelines, except that the final elution volume was 30 *µ*l instead of 100 *µ*l. DNA integrity was validated with a NanoDrop Spectrometer and PCR combined with electrophoresis of the Glutamine synthetase gene (glnA) as this gene is ubiquitous across microbial life (Supplementary Figure 1 and Supplementary Table 5). Adequate DNA concentrations for sequencing were reached by combining triplicate extractions for a total volume of 45 *µ*l and a concentration ≥ 50 ng/*µ*l (Supplementary Table 4). Positive and negative controls were used to verify the integrity of the PCR amplification. Two kit extraction blanks contained no trace of DNA and were not sequenced.

### Chemical analyses

Redox potential, pH, and dissolved O_2_ concentration profiles were measured at 100 *µ*M intervals in the field within an hour of collection, using Unisense (Aarhus, Denmark) microsensors connected to a Unisense Field Multimeter. Cores used for porewater chemistry analysis were sectioned in 1 cm intervals into 50 mL falcon tubes, followed by flushing of any headspace with ultra-high-purity nitrogen (N_2_) before capping. Sediment porewater was extracted following centrifugation at 4,000 rpm. The supernatant was then filtered through 0.45 *µ*m cellulose acetate filters into 15 ml tubes, and were frozen until analysis. Concentrations of nitrates and nitrites 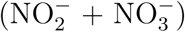, and ammonia (NH_3_), chloride (Cl^*−*^) were measured in the sediment porewater using a Lachat QuickChem 8500 FIA Ion Analyzer, while total dissolved phosphorus (TDP) and 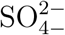 were measured in the sediment porewater using an ion chromatograph at the Biogeochemical Analytical Service Laboratory (Department of Biological Sciences, University of Alberta). However, TDP was removed from data analysis because insufficient porewater was collected to measure TDP at site C. The centrifuged sediments were retained and percentages of calcium carbonate (CaCO_3_) and organic carbon (OC) were estimated through loss on ignition (Heiri *et al.*, 2001).

The chemical features of the top 5 cm of the sediment cores were derived from measurements performed at 1 cm intervals throughout the cores. The geochemical properties of each sediment site were summarized using a Principle Component Analysis (PCA) and projections were clustered using Partitioning Around Medoids (Maechler *et al.*, 2019). The appropriate number of clusters was determined from silhouettes with the R package hopach (van der Laan and Pollard, 2003). The Dunn test (Dinno, 2017) was used to compare samples, controlling for multiple comparisons with the Benjamini-Hochberg adjustment.

### Sequencing and data processing

Metagenomic libraries were prepared and sequenced by Genome Quebec on an Illumina HiSeq 2500 platform (Illumina, San Diego, CA, USA; Supplementary Figure 2) on a paired-end 125 bp configuration using Illumina TruSeq LT adapters (read 1: AGATCGGAAGAGCACACGTCTGAACTCCAGTCAC, and read 2: AGATCGGAAGAGCGTCGTGTAGGGAAAGAGTGT). The DNA from the eight sites (five sediments, three soils) was sequenced, generating over 150 GB of data. Read count summaries were tracked throughout each step of the pipeline for quality control (Supplementary Figure 3). Low quality reads, adapters, unpaired reads, and low quality bases at the ends of reads were removed to generate quality controlled reads with Trimmomatic (v0.36) (Bolger *et al.*, 2014) using the following arguments: phred33, ILLUMINACLIP:TruSeq3-PE-2.fa:3:26:10, LEADING:3 TRAILING:3, SLIDINGWINDOW:4:20, MINLEN:36, CROP:120, HEAD-CROP:20, AVGQUAL:20. FASTQC (v0.11.8) (Andrews *et al.*, 2010) was then used to confirm that the Illumina adapters were removed and that trimmed sequence lengths were at least 90 bp in length with a Phred score of at least 33.

### Reconstruction of environmental genomes and annotation

To reconstruct environmental genomes, metagenomic quality-controlled reads from all samples were coassembled using Megahit (Li *et al.*, 2015) software with a k-mer size of 31 and “meta-large” setting (see Supplementary Table 6 for additional summary statistics). EukRep (West *et al.*, 2018) was used to remove any eukaryotic DNA from the contigs prior to the formation of an Anvio (v5) (Eren *et al.*, 2015) contig database. The contig database was generated by removing contigs under 1000 bp, and gene prediction was performed in the Anvio environment. Sequence coverage information was determined for each assembled scaffold by mapping reads from each sample to the assembled contig database using Bowtie2 (Langmead and Salzberg, 2012) with default settings. The resulting SAM files were sorted and converted to BAM files using samtools (v0.1.19) (Li *et al.*, 2009). Each BAM file was prepared for Anvio using the “anvi-init-bam” and contig database generated using “anvi-gen-contigs-database”. The contig database and BAM mapping files were further used as input for “anvi-profile”, which generated individual sample profiles for each contig over the minimum length of 2500 bp. These profiles were then combined using “anvi-merge” and summary statistics for abundance and coverage were generated with “anvi-summarise.” Automated binning was performed using CON-COCT (Alneberg *et al.*, 2014). Scaffolds were binned on the basis of GC content and differential coverage abundance patterns across all eight samples. Manual refinement was done using Anvio’s refine option (Supplementary Table 7). Kaiju (Menzel *et al.*, 2016) was used to classify taxonomy of the assembled contigs with “anvi-import-taxonomy-for-genes” and aided in the manual refinement process. Open reading frames were predicted with Prodigal (v2.6.3) (Hyatt *et al.*, 2010). Anvio’s custom Hidden Markov models were run, along with NCBIs COG (Tatusov *et al.*, 2003) annotation to identify protein-coding genes. PFAM (Finn *et al.*, 2015), TIGRFAM (Haft *et al.*, 2003), GO terms (Ashburner *et al.*, 2000), KEGG enzymes and pathways (Kanehisa *et al.*, 2015), and Metacyc pathways (Caspi *et al.*, 2007) were predicted with Interproscan (v5) (Jones *et al.*, 2014). These annotations were then combined with the Anvio database with “anvi-import-functions”.

Genome completeness and contamination were evaluated on the presence of a core set of genes using CheckM (v1.0.5) “lineage wf” (Supplementary Table 7) (Parks *et al.*, 2015). Only genomes that were at least 50% complete and with less than 10% contamination were further analysed – meeting the MIMAG standard for medium or high-quality genomes (Bowers *et al.*, 2017). All recovered genomes were used to calculate an average amino acid identity across all genomes using compareM (v0.0.23, function “aai wf”; https://github.com/dparks1134/CompareM) (Parks *et al.*, 2017). CheckM was used again to identify contigs that were not contained in any of the 300 high-quality genomes, that is those whose size ranges from 1000–2500 bp. As an attempt to “rescue” these unbinned contigs, an alternative binning algorithm MaxBin (v2.0) (Wu *et al.*, 2015) was employed. An additional 481 genomes were recovered, but were not included in further analysis as only 21 genomes were of average completion *>*65% (Supplementary Data 1: https://github.com/colbyga/hazen_metagenome_publication/blob/master/Supplemental_Data_1_maxbin2_unbinned_contigs_summary.csv).

### Phylogenetic placement of the MAGs

Phylogenetic analyses were performed using two different sets of marker genes from the Genome Taxonomy Database (GTDB): one for bacteria (120 marker genes) and one for archaea (122 marker genes), as previously been used to assign taxonomy to MAGs (Parks *et al.*, 2018). The marker genes were extracted from each genome by matching Pfam72 (v31) (Finn *et al.*, 2015) and TIGRFAMs73 (v15.0) (Haft *et al.*, 2003) annotations from GTDB (v86) (Parks *et al.*, 2018). Marker genes from each of the 300 genomes were translated using seqinr (Charif and Lobry, 2007), selecting the genetic code that returned no in-frame stop codon. As some genomes had multiple copies of a marker gene, duplicated copies were filtered out by keeping the most complete sequence. Marker genes that were missing from some genomes were replaced by indel (gap) characters, and their concatenated sequences were added those from the reference GTDB sequences. MUSCLE (v3.8.31) (Edgar, 2004) was employed to construct the alignment in R (v 3.5.1) (R Development Core Team, 2008). Archaeal sequences were removed from the bacterial alignment on the basis of results from CheckM (Parks *et al.*, 2015) and independently verified using a custom list of archaea specific marker genes. Alignments were then refined using trimAI (Capella-Gutiérrez *et al.*, 2009) and the “-gappyout” parameter. FastTree2 (Price *et al.*, 2010), recompiled with double precision to resolve short branch lengths, was used to infer maximum likelihood phylogenetic trees from protein sequence alignments under the WAG +G model (Whelan and Goldman, 2001; Aris-Brosou and Rodrigue, 2012, 2019). The archaeal tree was rooted with Euryarchaeota and the bacterial tree was rooted with Patescibacteria using APE (Paradis *et al.*, 2004). Trees were visualized and colored by phylum with ggtree (Yu *et al.*, 2017).

### Community composition of the MAGs

To determine the relative abundance of each genome in the eight samples, sample-specific genome abundances were normalized by sequencing depth [(reads mapped to a genome) / (total number of reads mapped)], making comparisons across samples possible. Genome abundances were generated using the CheckM “profile” function (Parks *et al.*, 2015). To determine the average abundance of major taxonomic groups across sites (determined by the phylogenetic analysis described above), the abundances for genomes from the same taxonomic group were summed and visualized using phyloseq (McMurdie and Holmes, 2013) (usually at the phylum level, unless otherwise stated). These same abundance values were the basis for a community composition analysis. The *t*-SNE plots were constructed by assigning each genome to a site based on where it was most abundant using Rtsne (Krijthe *et al.*, 2018).

### Metabolic potential of the MAGs

To analyze functional marker genes in the metagenomes, we used a custom database of reference proteins sequences (COG, PFAM, TIGRFAM, KEGG) based on the marker genes used in other studies (Anantharaman *et al.*, 2016; Dombrowski *et al.*, 2018) (Supplementary Data Files on GitHub). Pathways were also predicted using MinPath (Ye and Doak, 2009) to map all identified KEGG enzymes to the most parsimonious MetaCyc pathways (Caspi *et al.*, 2007). As these MAGs were incomplete, some genes in pathways may be absent. MinPath presented only parsimonious pathways represented by multiple genes. As most genomes were present even at low abundances across all sites, a cut-off value of ≤ 0.25 (on a −*log*_10_ scale) was set for a genome to be included in the functional analyses at any site, so that only the most abundant genomes for each site were considered. We aggregated marker genes and pathways by function, summarizing the results by phyla, except for Proteobacteria that was separated by class. We further grouped all taxa together at each site to test for significant differences in major nutrient cycling processes (carbon, nitrogen, and sulphur) among sites using a hierarchical clustering; significance was derived from the Approximately Unbiased bootstrap (Suzuki and Shimodaira, 2006) and Fisher’s exact test.

## Results

### Characterization of the physical and geochemical environments

We first characterized how geochemical properties of the sediments varied along and between the two transects. Sediment samples from these five sites clustered into four distinct geochemical groups (Figure 1B) that reflect spatial variability in glacial runoff, the primary hydrological input to the lake. Indeed, PC1 explained 43% of the total variance (*σ*^2^), and differentiated the L and high H runoff transects, while PC2 (29.9%) separated each transect according to their depth.

Along PC1, higher concentrations of ammonia (NH_3_) and sulfate 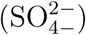 in the pore-waters, and a greater percentage of calcium carbonate in the sediments, were present in the H transect. However, higher concentrations of dioxygen (O_2_), nitrates / nitrites 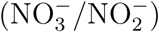, and greater redox potential were present in the L transect and the control (C) sites. Along PC2, sediment organic carbon (OC), and porewater pH and Cl^*−*^, were more determinant when discriminating between the shallow (L1 and H1) and deep (L2 and H2) sites of both transects (Supplementary Figures 4-5). Rather than grouping spatially with the H transect, the C sites were most chemically similar to L1 (Figure 1C, Supplementary Figure 6). The shallow sites were not significantly different from each other in pH or OC concentrations, but were both significantly different from the deeper sites suggesting that although most chemical features were similar within each transect, some features might still be influenced by their spatial proximity to the shoreline or depth of the overlying water column (Figure 1C).

### Contrasting low *vs.* high runoff transects revealed a decrease in biodiversity

With such a clear geochemical separation of the transects along PC1 (43% of *σ*^2^) and significant spatial contrasts (Figure 1C), we had the right context to evaluate the influence of runoff gradients on sediment microbial diversity. We assembled a total of 300 (290 bacterial and 10 archaeal) MAGs that were *>*50% complete and with *<*10% contamination (Supplementary Tables 6-7). By constructing phylogenetic trees for Bacteria and Archaea, we noted that while most major phyla were represented in the MAGs, no Firmicutes and only a small number of Archaea were identified (Figure 2). In contrast, Gammaproteobacteria (*n* = 50), Actinobacteria (*n* = 31), Alphaprobacteria (*n* = 24), Chloroflexoata (*n* = 30), Planctomycetota (*n* = 24), and Acidobacteriota (*n* = 19) were the most commonly recovered taxa across the entire watershed. Uncultured phyla comprised ∼11% of reconstructed MAGs, including representatives from multiple taxa: Eisenbacteria (*n* = 12), Patescibacteria (*n* = 9), Omnitrophica (*n* = 5), KSB1 (*n* = 1), Armatimonadota (*n* = 1), Lindowbacteria (*n* = 1), USBP1 (*n* = 1), UBP10 (*n* = 1), and Zixibacteria (*n* = 1).

**Figure 2.**
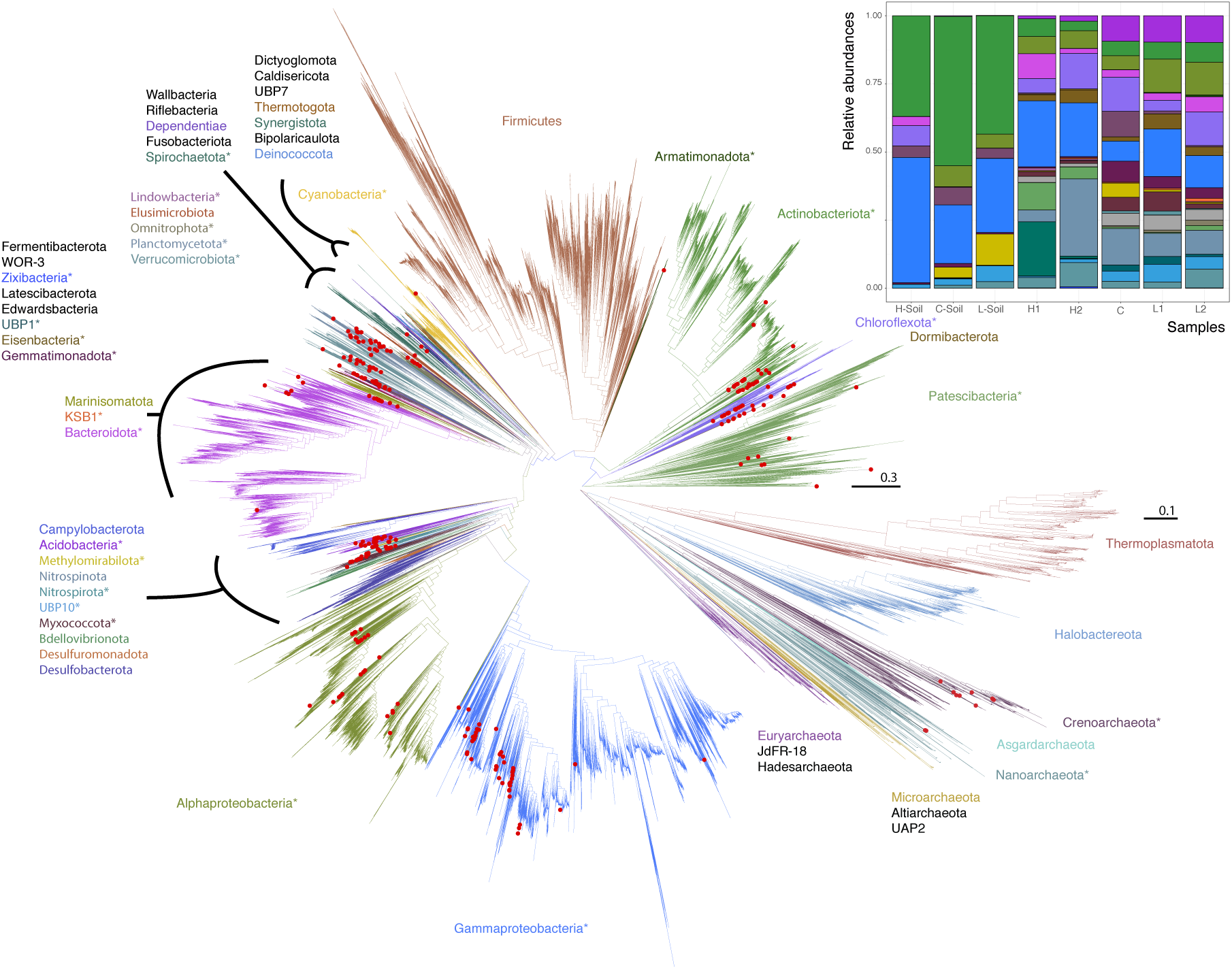
Maximum likelihood phylogenetic trees of Lake Hazen genomes based on 120 concatenated bacteria and 122 concatenated archaea protein-coding genes. Red Dots: Lake Hazen genomes. Asterisks (*) indicate phyla that contain Lake Hazen genomes. Bacteria tree is rooted with Patescibacteria and Archaea tree is rooted with Euryarchaeota. See GitHub account for full taxonomy tree files and for original tree files (Supplemental Data File 2 and 3). Inset shows MAG abundance across sites, in the 300 high quality genomes for each sample normalized to 100%.

However, these MAGs were not evenly distributed across all sites (Figure 2, inset; Supplementary Figure 7). To quantify this uneven distribution, we determined the site where each genome was most abundant. Based solely on this information, we performed an unsupervised clustering (*t*-SNE), and found that the directions defined by sediment-laden water flowing from the shallow to the deep site within each transect in the projection space were almost orthogonal between transects (see arrows in Figure 3). This orthogonality suggests that transitioning from the L to the H transect could lead to a dramatic shift in microbial communities.

**Figure 3.**
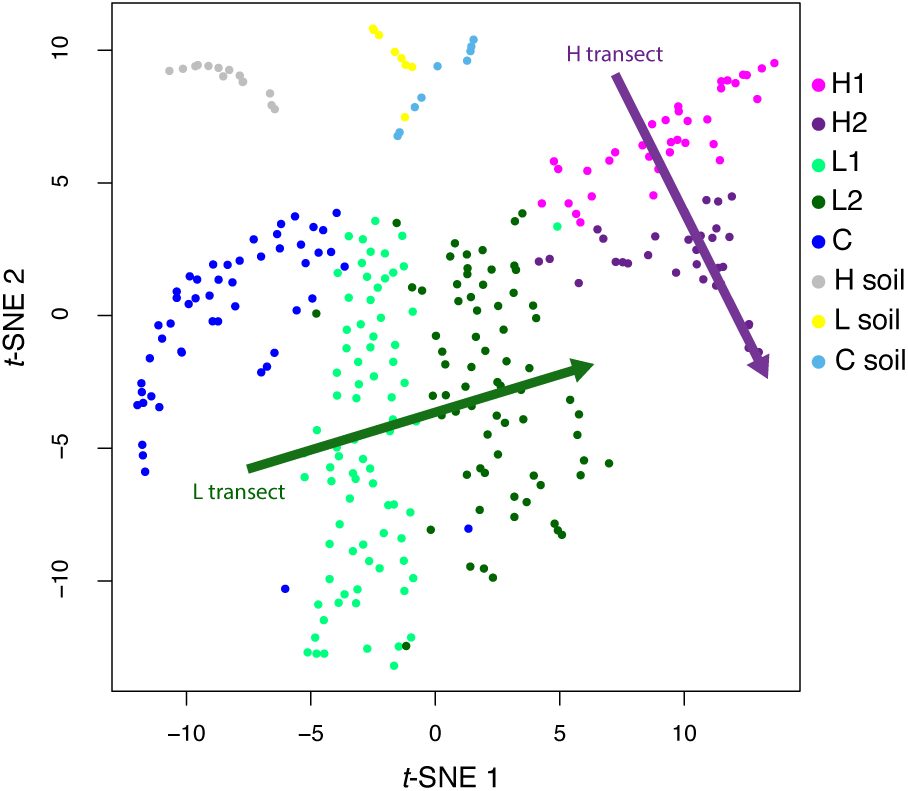
*t*-SNE analysis of genome abundance for each sediment sample. Each of the 300 shown genomes was assigned to the sample where it has the greatest abundance. Shaded arrows display the approximate direction of water flow, from upstream to downstream, for the high (green) and low (purple) transects.

To assess the significance of these shifts at the phylum level, we calculated the relative proportions of each of the reconstructed 300 MAGs at each site, and tallied these numbers by phylum, over the 43 phyla represented in our data. We did this along each transect – essentially pooling sites H1/H2 together to represent the H transect, and doing the same for sites L1/L2 (the L transect), while keeping proportions for the S and C sites separate. Hierarchical clustering on this table of MAGs proportions by phyla *vs.* sites showed a divergence from the L to H transects (following the (((L,C),H),S) clustering pattern; Figure 4A, inset), confirming the clear contrast between the two transects in terms of taxa proportions (see Figure 3). To test if these taxa proportions tended to increase or decrease when transitioning from L to H along the (((L,C),H),S) clustering pattern, we fitted linear models (ANOVA) regressing the proportions of each of the 43 phyla against sites, ordered as per their hierarchical clustering (L→C→H→S). Essentially, we regressed a single data point for each of the four classes (L, C, H, and S), so that *P* -values could not be obtained, but slope could be estimated (Figure 4A). Strikingly, most of these slopes were negative (binomial test: *P* = 7.8 × 10^*−*8^), demonstrating a significant decrease in diversity at the phylum level as one goes from low to high runoff regimes.

**Figure 4.**
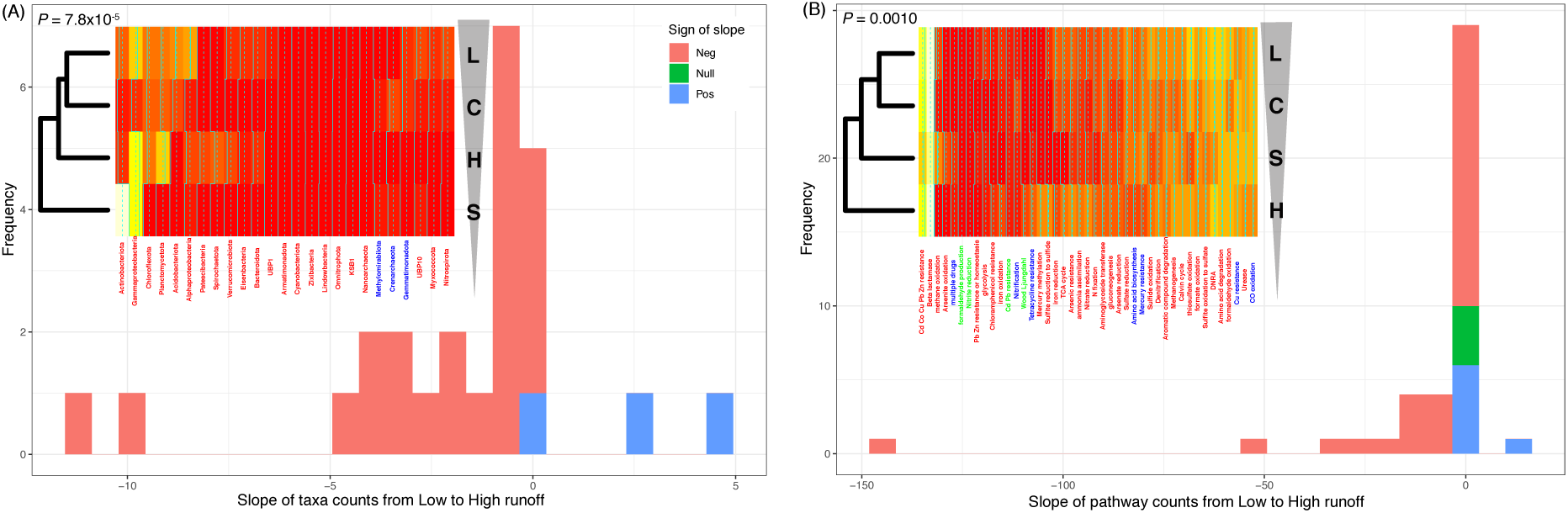
Transition from low to high runoff leads to a decrease in diversity. (A) Distribution of the slopes of taxonomic counts as a function of sites. (B) Distribution of the slopes of pathway counts as a function of sites. In both cases, counts were aggregated by location types (L [Low], C [Control], S [Soil], and H [High] sites), and linear models (ANOVA) were fitted to estimate the slope of each regression. Insets: heatmap representations of count tables; leftmost dendrograms show how the location types cluster, transitioning from L to H runoffs (vertical triangle pointing down). *P* -values: one-sided binomial test for enrichment in negative slopes.

An NMDS ordination allowed us to detect the geochemical features associated with this shift in microbial communities (Supplementary Figure 8). In the sediments, NH_3_ concentrations (*P* = 0.03), 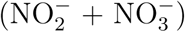 concentrations (*P* = 0.03), and redox potential (*P* = 0.03) were significant in determining the distribution of MAGs (permutation test: *P <* 0.05). We further observed that the sites with the greatest diversity (L/C sites) were also those with the greatest redox potential, and O2 and 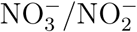 concentrations. Sites with the lowest microbial diversity (H sites), contained greater NH_3_ and 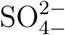 concentrations, and lower redox potential. In addition to gradients shaped by the interplay between microbial metabolism and local geochemical constraints, the physical disturbances associated with high sedimentation rates also likely contributed to the homogenization of the microbial community structure; however, we cannot quantify the relative importance of each of these processes here.

### Contrasting low *vs.* high runoff transects also revealed a loss of functional potential

To assess the functional implications of this decrease of biodiversity, we assigned metabolic functions and pathways to proteins in each MAG. We focused on genes and pathways involved in key elements, targeting carbon, nitrogen, and sulfur cycling (Supplementary Figures 9-10). Only the most abundant genomes per site were reported within each phylum (Supplementary Figure 11), allowing us to compute the proportions of functions and pathways in each of the 43 phyla present in reconstructed MAGs across the hydrological regimes. Their hierarchical clustering (Supplementary Figures 12-14) led to a picture consistent with the ones derived from both geochemical (Figure 1) and taxonomic abundances (Figure 4A). Indeed, the two transects were again clearly separated (clustering pattern (((L,C),S),H); Figure 4B, inset), and fitting linear models regressing function/pathway proportions against sites showed that, again, most of these slopes were negative (binomial test: *P* = 0.0010). Forcing the same site ordering as for the taxonomic abundances (L→C→H→S as in Figure 4A, inset) led to similar results (binomial test: *P* = 7.8×10^*−*5^), demonstrating a significant decrease in metabolic diversity when going from the L to the H transect.

More specifically, we found that marker genes whose product is implicated in carbon and sulfur metabolisms significantly decreased when going from the L to H, while nitrogen metabolism was unaffected (Supplementary Table 8; see Supplementary Text for details). When considering the individual functions present or absent across the transects, we noted that most oxidative pathways (CO, methane, formaldehyde, sulfide, sulfite) appeared less common in the H transect (Supplementary Figure 9), corresponding to lower oxygen concentrations and constraints on aerobic metabolism. Furthermore, while most carbon fixation processes were shared between the two transects, carbon oxidation and reduction reactions regulated through Wood-Ljungdahl pathway were only observed in the H transect, where sedimentary conditions were anoxic throughout the first 5cm (Supplementary Figures 4-5), consistent with a more reductive environment.

## Discussion

Even if Arctic microbial communities are changing rapidly (Hultman *et al.*, 2015), there is still a dearth of long-term time series observations. To address this point, we used Lake Hazens spatial geochemical heterogeneity to evaluate the structural and functional response of lake sediment microbial communities to varying runoff, already shown to increase in this warming High Arctic environment (Lehnherr *et al.*, 2018). Such an approach can reasonably be interpreted from the lens of a space-for-time design, which assumes that spatial and temporal variations are not only equivalent (Blois *et al.*, 2013; Lester *et al.*, 2014), but also stationary (Damgaard, 2019). Whether this latter condition is met cannot be known, but in the absence of any time-series documenting the effect of climate change on lake sediment microbial communities in the High Arctic, the space-for-time design becomes a convenience, if not a necessity (Pickett, 1989).

Using metagenomics along two transects experiencing heterogeneous runoff conditions, we presented evidence that climate change, as it drives increasing runoff and sediment loading to glacial lakes, will likely lead to a decrease in both diversity and functional potential of the dominant microbial communities residing in lake sediments. Note that we specifically focused here on the dominant microbes, that is those for which we could reconstruct the MAGs, in order to (i) have a phylogenetic placement of the corresponding organisms based on a large number of marker genes (Figure 2), rather than partial 16S rRNA gene sequences as usually done (Ruuskanen *et al.*, 2018b), and (ii) be able to predict almost complete functional pathways for each of these organisms to test the impact of a change of runoff (Figure 4), rather than inferring function from taxonomic affiliation (Ruuskanen *et al.*, 2018b).

Such a decrease in taxonomic and functional diversity may not be unique to Lake Hazen, where rising temperatures have resulted in increasing glacial melt and associated runoff. Although such a pattern was not observed in other regions of the globe where runoff is predicted to decrease (Huss and Hock, 2018; Pierre *et al.*, 2019), our finding are likely to apply to other, smaller, glacierized watersheds typical of high latitudes or altitudes. Indeed, at least in the Arctic, freshwater discharge is broadly expected to increase with increasing temperatures and precipitation loadings (Peterson *et al.*, 2002; Rawlins *et al.*, 2010; Bring *et al.*, 2016). It would thus be immensely valuable to conduct similar studies, replicating where appropriate a similar space-for-time design, at other lakes throughout the world. Additional sampling efforts should carefully consider the spatial heterogeneity of runoff regimes leading to divergent sedimentation rates (Supplementary Table 2), limiting our ability to make temporal predictions.

Despite lacking geochemical measurements for the soil samples, we found that the microbial communities in the sediments at the high runoff sites clustered most frequently with those in the soil sites (Figure 4), highlighting a connection between terrestrial and aquatic sediment communities as a function of the runoff volume, consistent with previous findings (Comte *et al.*, 2018; Ruiz-González *et al.*, 2015). Unsurprisingly, as the soil is likely a source of nutrients (*e.g.*, DOC) and organic and inorganic particles, we would expect increased runoff to the aquatic ecosystems to alter microbial community structure (Le *et al.*, 2016). Some of these structural changes may then alter the functional capacity to metabolize carbon, nitrogen, sulfur compounds and process toxins such as metals and antibiotics (Supplementary Figure 9). A more experimentally-driven approach, based for instance on *in situ* incubation and geochemical tracers, would have been necessary to quantify such an interplay between microbial metabolism and geochemical features. Yet, as sediments and nutrients are mostly deposited during the summer melt months, it can be expected that lake sediments record microbe-driven seasonal changes in their geochemistry. Indeed, high glacial runoff is known to bring dense, oxygenated river waters with OC directly to the bottom of the lake (Pierre *et al.*, 2019), stimulating aerobic microbial activity. As a result, the geochemistry recorded along the high runoff transect may first reflect a period of greater microbial metabolism, which may actually exceed those in temperate systems (Probst *et al.*, 2018), eventually followed by low oxygen, low redox, and high NH_3_ conditions observed here (Figure 1) as oxygen is depleted and anaerobic metabolisms allowed to proceed.

At a larger temporal scale, a key question that arises from these results is how changes in hydrological regimes will alter the evolutionary dynamics of microbial communities in lake sediments. Niche differentiation, where the coexistence of ecological opportunities can facilitate species diversification, may explain why sediments along the low runoff transect hosts a more diverse microbial community than sediments along the high runoff transect (Cordero and Polz, 2014). Presently, climate change is predicted to increase runoff in this High Arctic environment (Lehnherr *et al.*, 2018), and we found evidence suggesting that the increased runoff homogenizes community structure. This can be expected to disrupt niche differentiation, and hence to reduce the overall and long-term metabolic capacity in lake sediments. It is currently hard to predict the future microbial ecology of these systems. On the one hand, climate change may diminish species diversification, and lead to highly specialized microbial communities adapted to a homogeneous ecological niche characterized by low oxygen, low redox, and high NH_3_ concentrations. On the other hand, the seasonal and rapid changes in redox conditions, predicted to follow the strong but punctual input of oxygen and nutrients during springtime may allow for the development of a short-lived community that eluded our sampling and analysis.

The rapid changes that affect Lake Hazen’s watershed in response to climate warming were already known to directly alter its hydrological regime. Here we further provide evidence that a combination of increasing runoff and changing geochemical conditions are associated with the reduced diversity and metabolic potential of its dominant microbial communities. While longitudinal studies are needed to confirm these patterns, it is still unclear how such losses in biodiversity and metabolic potential in Arctic ecosystems will impact key biogeochemical cycles, potentially creating feedback loops of uncertain direction and magnitude.

## Acknowledgements

This study was made possible through a collaborative effort undertaken by Igor Lehnherr, Stephanie Varty, Victoria Wisniewski (University of Toronto, Mississauga), Charles Talbot (Environment and Climate Change Canada), and Maria Cavaco (University of Alberta). We thank Linda Bonen, Marina Cvetkovska, Manon Ragonnet and Alex Wong for comments and discussions. Funding support was provided by the Natural Science and Engineering Research Council of Canada (VSL, AJP, SAB), ArcticNet Network Centre of Excellence (VSL, AJP), and the Polar Continental Shelf Program (VSL) in Resolute, Nunavut, which provided logistical and financial support.

## Author contributions

G.C. and V.S.L. performed sampling, whereas G.C. conducted laboratory analyses. G.C. and S.A.B. performed data analyses. G.C., S.A.B., V.S.L., and A.J.P. designed the study and wrote the manuscript. V.S.L. conducted the microsensor profiles and porewater extractions. G.C., S.A.B., A.J.P., M.R., K.S.P., and V.S.L. reviewed the manuscript.

