## Supplementary Material for "Climate change negatively impacts dominant microbes in the sediments of a High Arctic lake"

### Supporting Text

**Nutrient cycles affected when transitioning from low to high.** Overall, markers of carbon and sulphur metabolism significantly decreased when transitioning from the L to H sites, even if nitrogen metabolism was not (Tab. S8). Most carbon pathways, such as carbon fixation through the Calvin-Benson-Bassham (CBB) pathway, as well as the capacity for simple carbon metabolism, were shared across all runoff regimes. In contrast, carbon oxidation and reduction reactions regulated through Wood-Ljungdahl pathway were only observed in the H sites, where sedimentary conditions were anoxic throughout the first 5 cm (Figs. S4, S9). Here, Spirochaetota were likely performing anaerobic respiration and carbon fixation producing acetate as an end product. Methanogenesis pathways were present across all sites, but notably, methane oxidation pathways were absent from high runoff sites, where oxygen is limited.

Greater concentrations of ammonia in the high runoff regimes may suggest that N-containing organic matter was mineralised through ammonification (Fig. S8). In the high runoff regime, there was both an absence of nitrification and a greater presence of markers for ammonia assimilation. Markers for dissimilarity nitrate reduction (DNRA) were present in multiple genomes across all runoff regimes (Fig. S8). In contrast, urease markers were found more abundantly in low runoff regimes, where ammonia concentration was lower (Fig. S5). The functional ability of microbes to cycle sulphur between oxidised and reduced forms was significantly different between the high and low runoff regimes (Tab. S8). In the high runoff regime, Gammaproteobactiera were the only organisms with the metabolic capacity to expansively utilise sulphur, performing sulphide oxidation and thiosulphate reduction. Whereas, sulphate reduction was predominantly found in the soil, control, and low runoff regimes.

Aside from nutrient cycling, we also assessed the capacity of microbial communities to process metals and antibiotics. Metal resistance and cycling was mostly ubiquitous throughout all of the sites, regardless of runoff. Methyl mercury production, identified by the presence of both hgcA and hgcB genes [1], was only implicated in the high runoff sites, in Spirocheatoa and Chloroflexoata. However, genes conferring mercury resistance involved in the conversion of inorganic $Hg^{II}$ to the less toxic $Hg^0$ – were evenly distributed throughout the sites. There was a broad presence of metal tolerance that was indicated by genetic determinants related to heavy metal resistance of cadmium, cobalt, copper, lead and zinc (Fig. S9). Furthermore, antimicrobial resistance genes specifically $\beta$-lactamases, were ubiquitous across all genomes at all sites. Finally, we found that while amino acids were readily synthesized and degraded by most organisms (Fig. S9), the degradation of polycyclic aromatic compounds appeared to be least prevalent in the high runoff sites.

**Sites clustered by runoff regime.** To identify potential drivers of this reduction in diversity when going from the L to the H transects, we bi-clustered genomes by their normalised abundances (on a -log10 scale to reduce skew) and by sample site (Fig. S12), and found that sites clustered following a similar pattern to geochemical features (see Fig. 1B), with H sites grouping separately from L sites (Fig. 3A). The normalised abundances of MAGs showed no strong phylogeographic pattern, in that we did not observe an assemblage of MAGs solely representative of a given site (Fig. S13). In spite of this absence of phylogeographic pattern, the tanglegram suggests that the beta diversity of highly abundant MAGs in the L/C sites was greater than at the H sites (see the distribution of green lines connecting the phylogenetic and clustered trees in Tab. S13). This difference in diversity between samples is further supported by an NDMS ordination (Fig. S14) and a PERMANOVA test on a PCoA ordination (Fig. S14).

### Supporting Figures

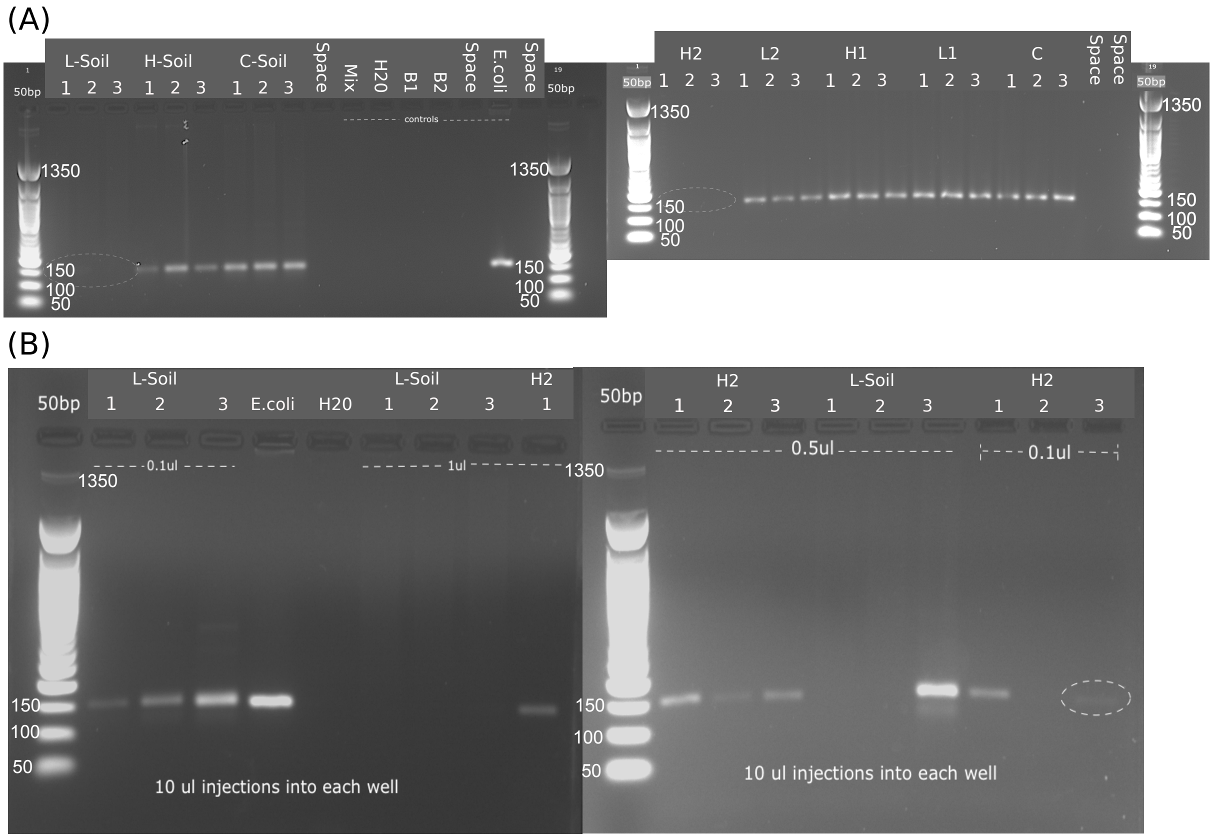


**Supporting Figure 1.** Gel electrophoresis images of the *glnA* gene for each sample extracted in triplicate. (A) Initial gel results using 1 $\mu$l of DNA for each PCR reaction. Every sample contains *glnA* except L-Soil and H2. (B) Repeated PCR reaction for L-Soil and H2. Diluted DNA concentrations for PCR from 1 $\mu$l to 0.1 $\mu$l and 0.5 $\mu$l.

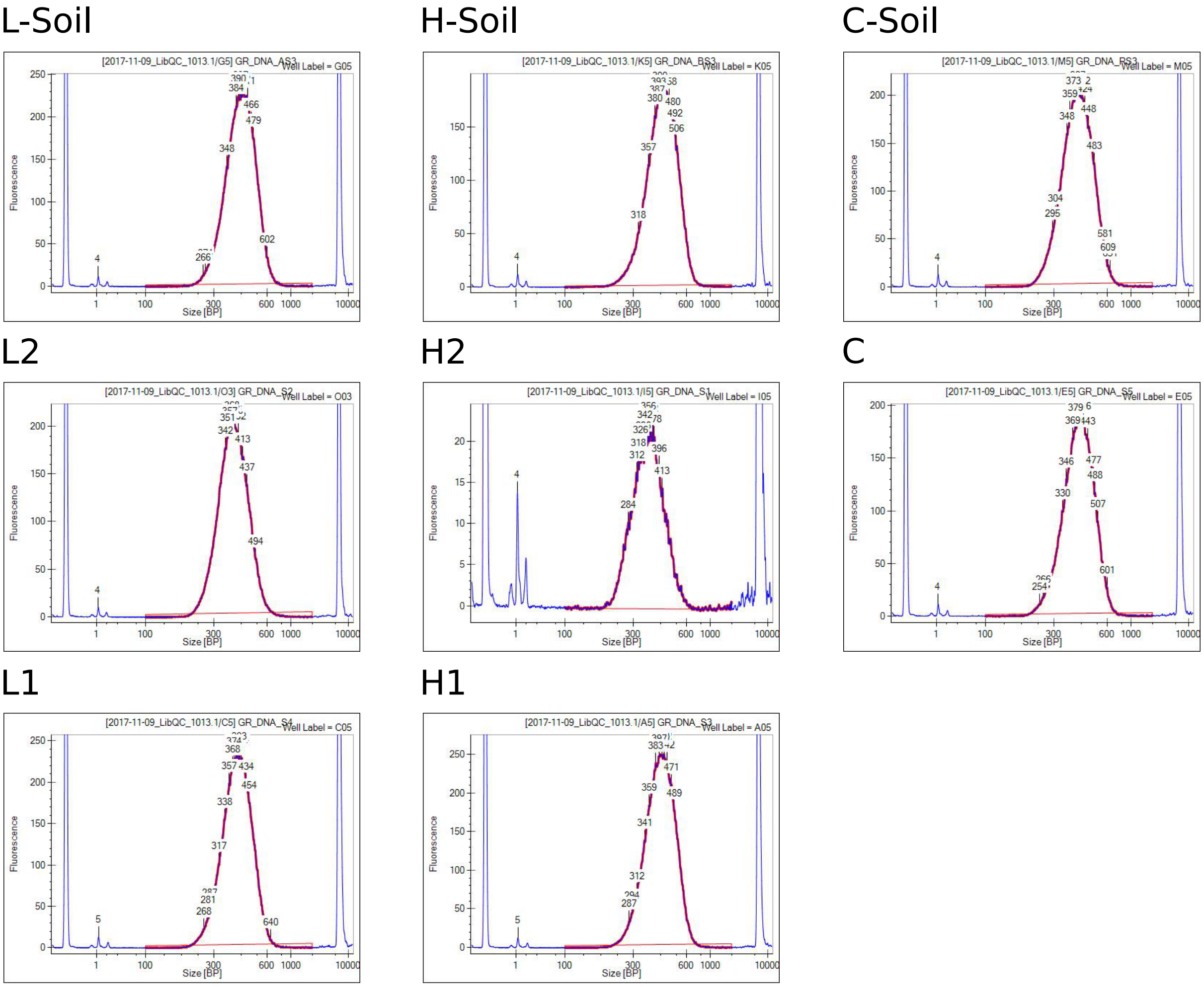


**Supporting Figure 2.** Illumina HiSeq DNA sequencing library validation from an Agilent Technology 2100 Bioanalyzer. All libraries were constructed using Illumina TruSeq DNA PCR-Free Library Prep. The library results for each of the eight samples is presented and labelled above the panel: L-Soil, H-Soil, C-Soil, L2, H2, C, L1, H1. Peaks between 300-600 bp indicate the presence of DNA. Peaks below 1 bp and above 1000 bp are internal standards.

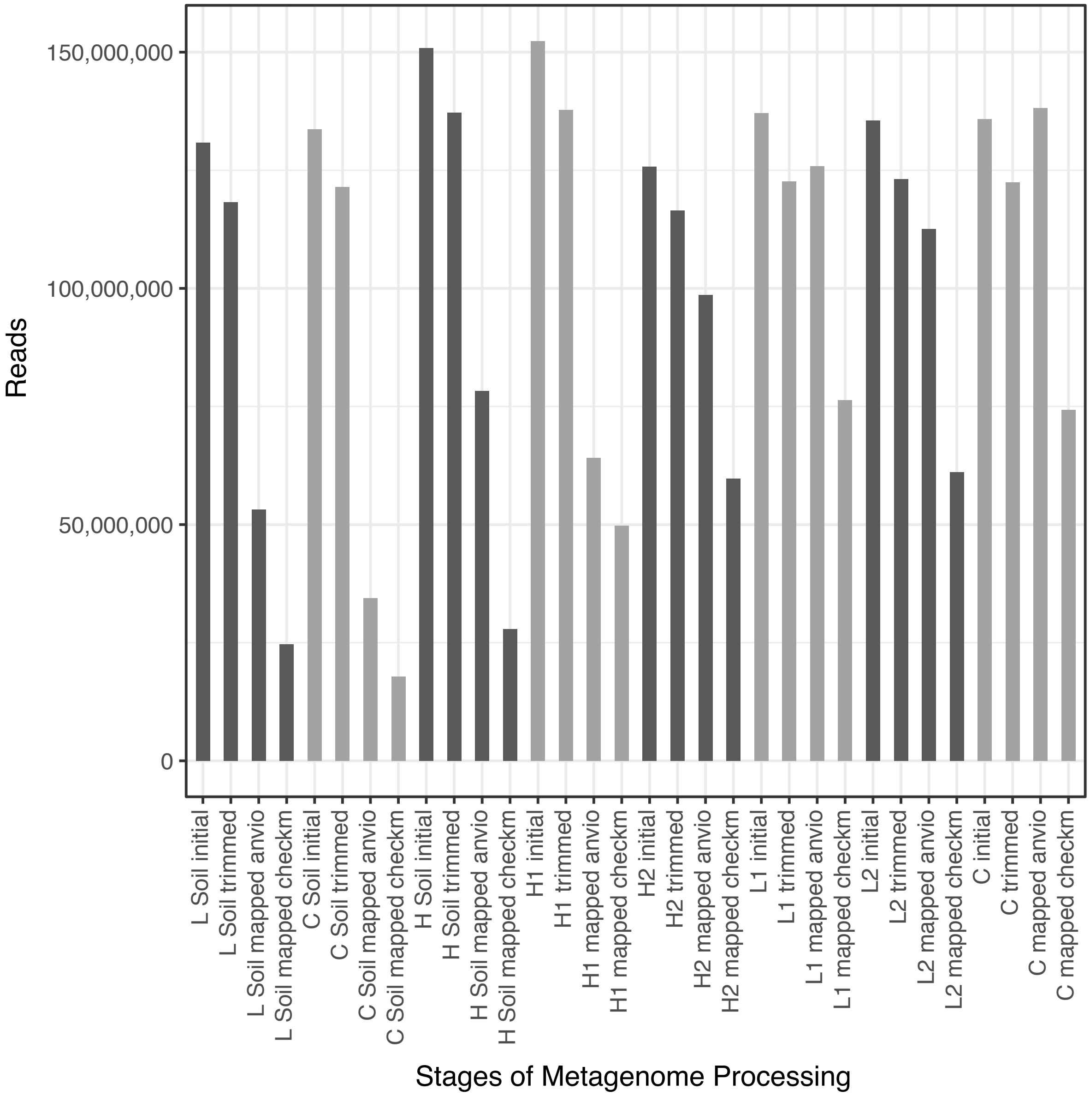


**Supporting Figure 3.** Number of reads throughout analysis. Reads are successively decreasing, except for the reads mapped to the Anvio contig database, as each read can map to more than one contig and be counted more than once. The reads mapped using checkM include only reads that map to the 300 high quality reconstructed genomes, opposed to all reconstructed genomes in the Anvio mapping.

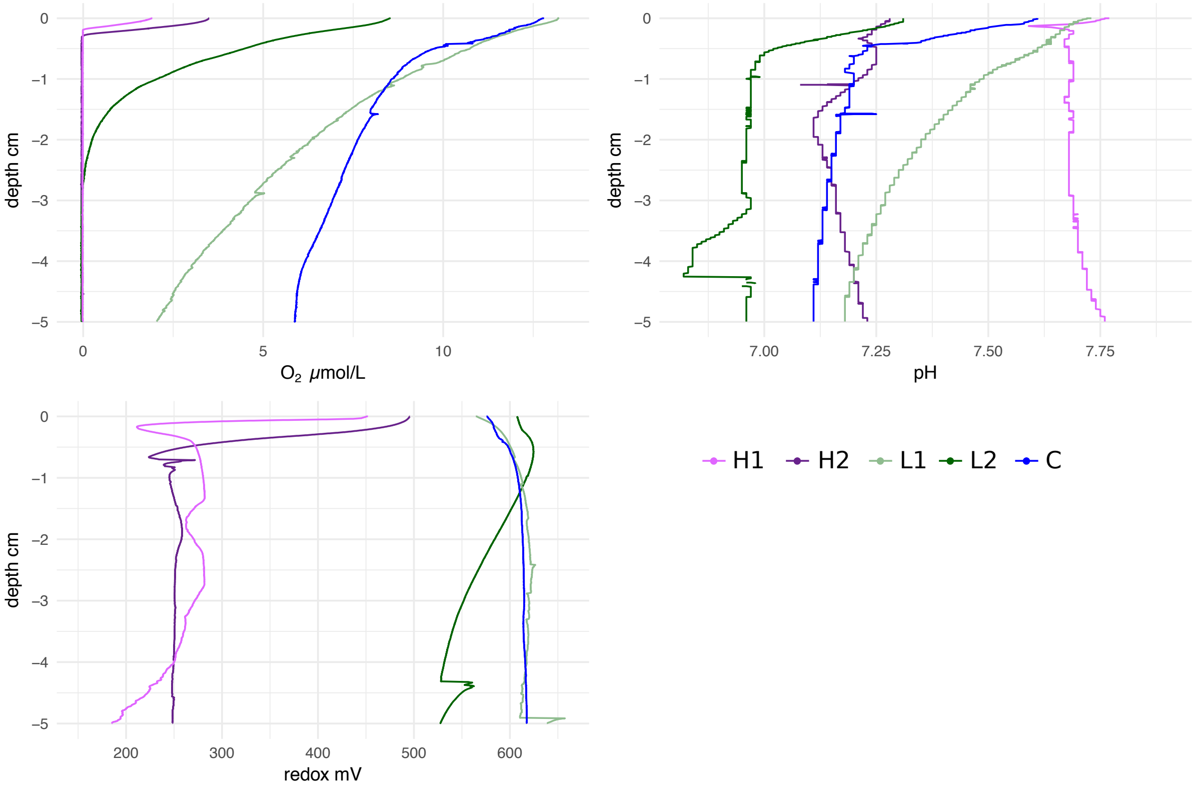


**Supporting Figure 4.** Sediment microprobe profiles for oxygen ($O_2$), pH, and redox measured in 100 $\mu$m intervals. Legend (bottom right) indicates the colour of each sediment site in the profiles.

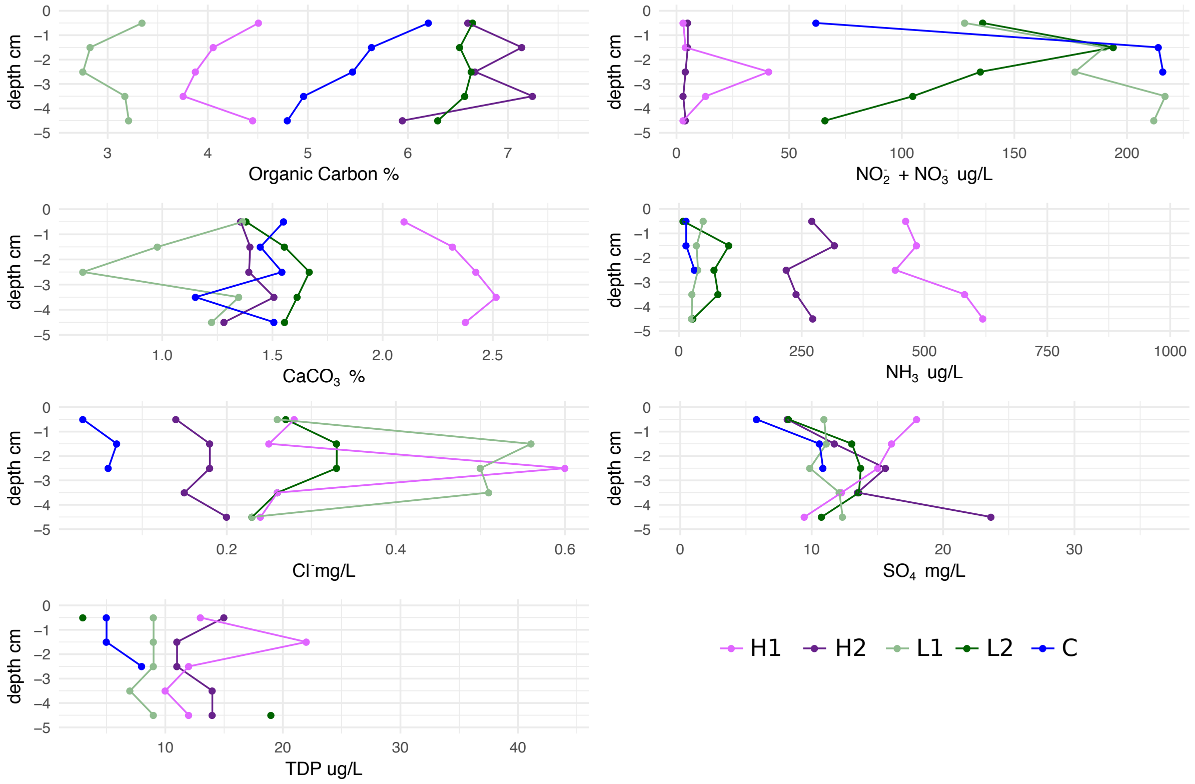


**Supporting Figure 5.** Sediment porewater profiles for organic carbon, nitrite and nitrates, calcium carbonate, ammonia, chlorine, sulphate, and total dissolved phosphorus (TDP). TDP was removed when producing PCA and boxplots in Figure 1 because of incomplete measurements in C and L2. Legend (bottom right) indicates the colour of each sediment site in the profiles.

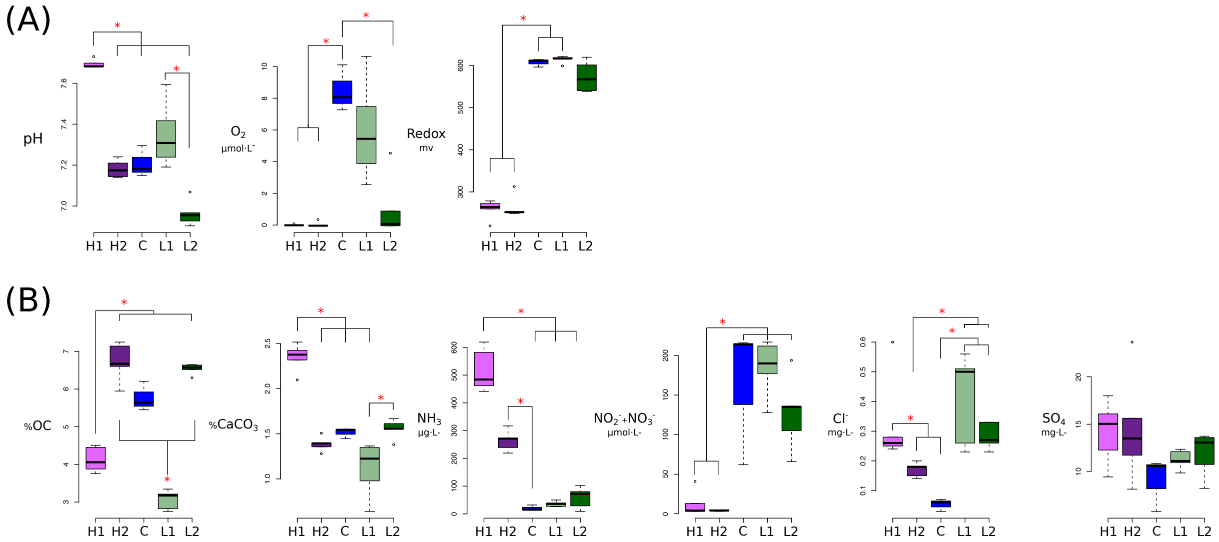


**Supporting Figure 6.** Distribution of all chemical features for sediment sites. Includes chlorine and sulphate measurements absent from Figure 1. Branches and asterisks indicate significant differences between sites $P < 0.025$ (Dunn Test). If branches form a dichotomy or trichotomy, the interactions within that group is not significant. (A) Microprobe measurements collected at every 100 $\mu$m. (B) Porewater measurements collected from bulk 1 cm intervals.

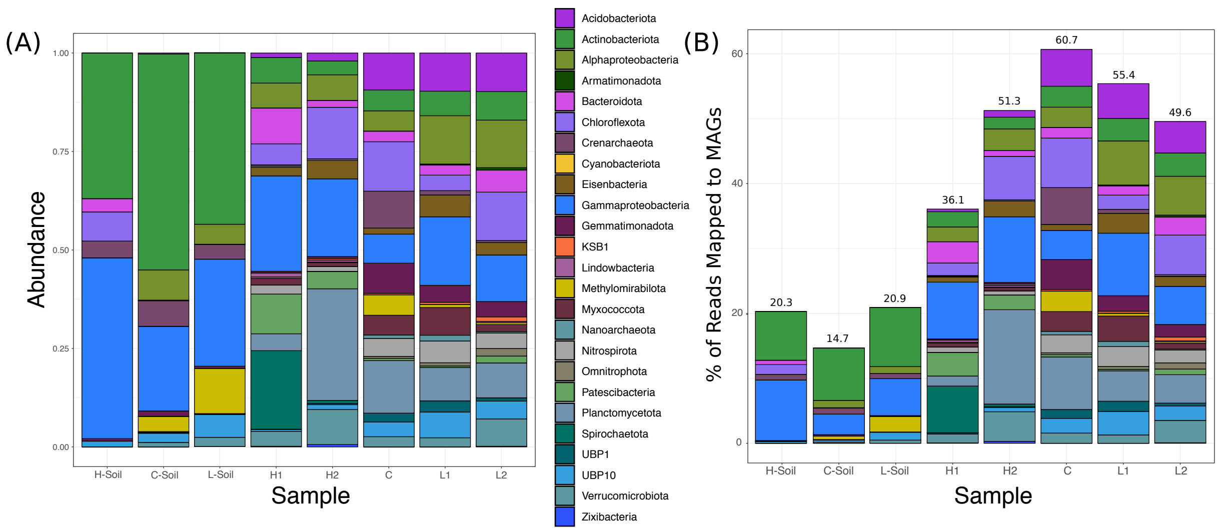


**Supporting Figure 7.** Reconstructed genome abundance across sites. (A) Amount of reads mapped to the 300 high quality genomes for each sample normalised to 100%. Only reads that were mapped to genomes are shown and all unmapped reads have been excluded. (B) Amount of reads mapped to the 300 high quality genomes for each sample.

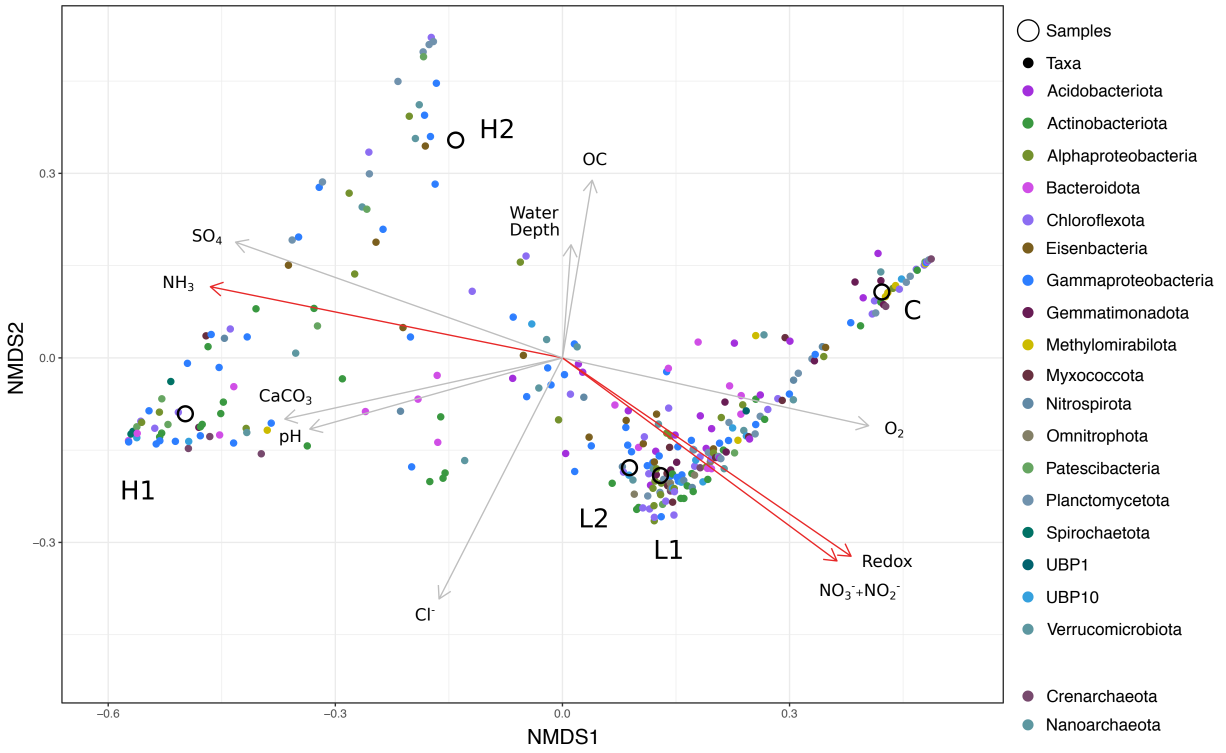


**Supporting Figure 8.** NMDS analysis of genomes from 20 most abundant phyla in sediment sites. Physical and chemical vectors are fitted to data. Red vectors are significant (permutation test: $P < 0.05$).

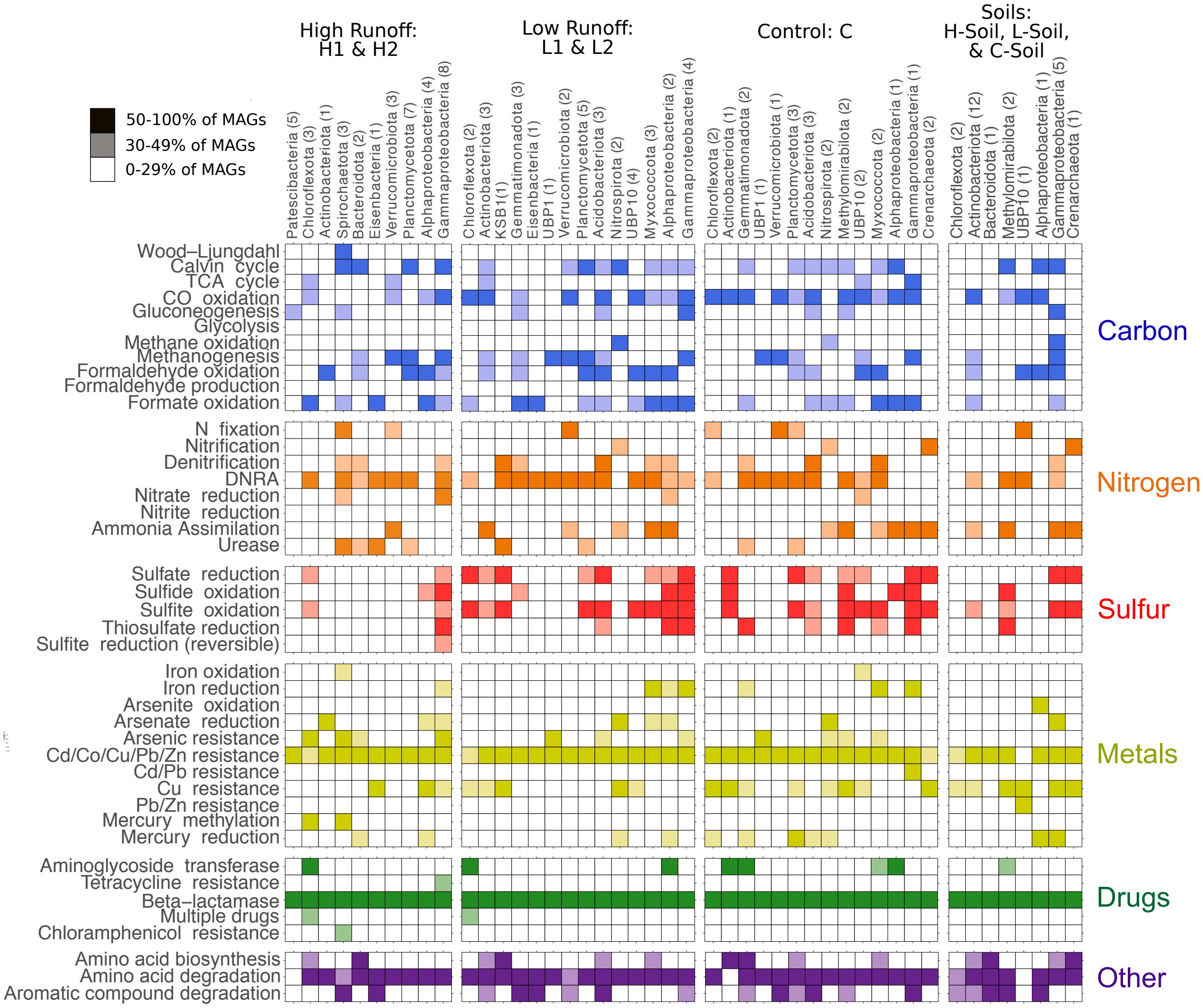


**Supporting Figure 9.** Metabolic capacity of genomes separated by hydrological regime. Genomes are only considered to contribute to a site if 0.56% ($-\log_{10} \leq 0.25$) of reads per sample mapped to a genome. Presence of core metabolic genes involved in carbon metabolism, nitrogen metabolism, sulphur metabolism, metal cycling, antibiotic resistance, and other metabolism are shown. Number of genomes for each taxa are shown in parentheses. Blank: genes predicting function are absent or in low abundance. Shaded colours: genes predicting function are present in 30-50% of genomes per phylogenetic group. Dark colours: genes predicting function are present in >50% of genomes per phylogenetic group.

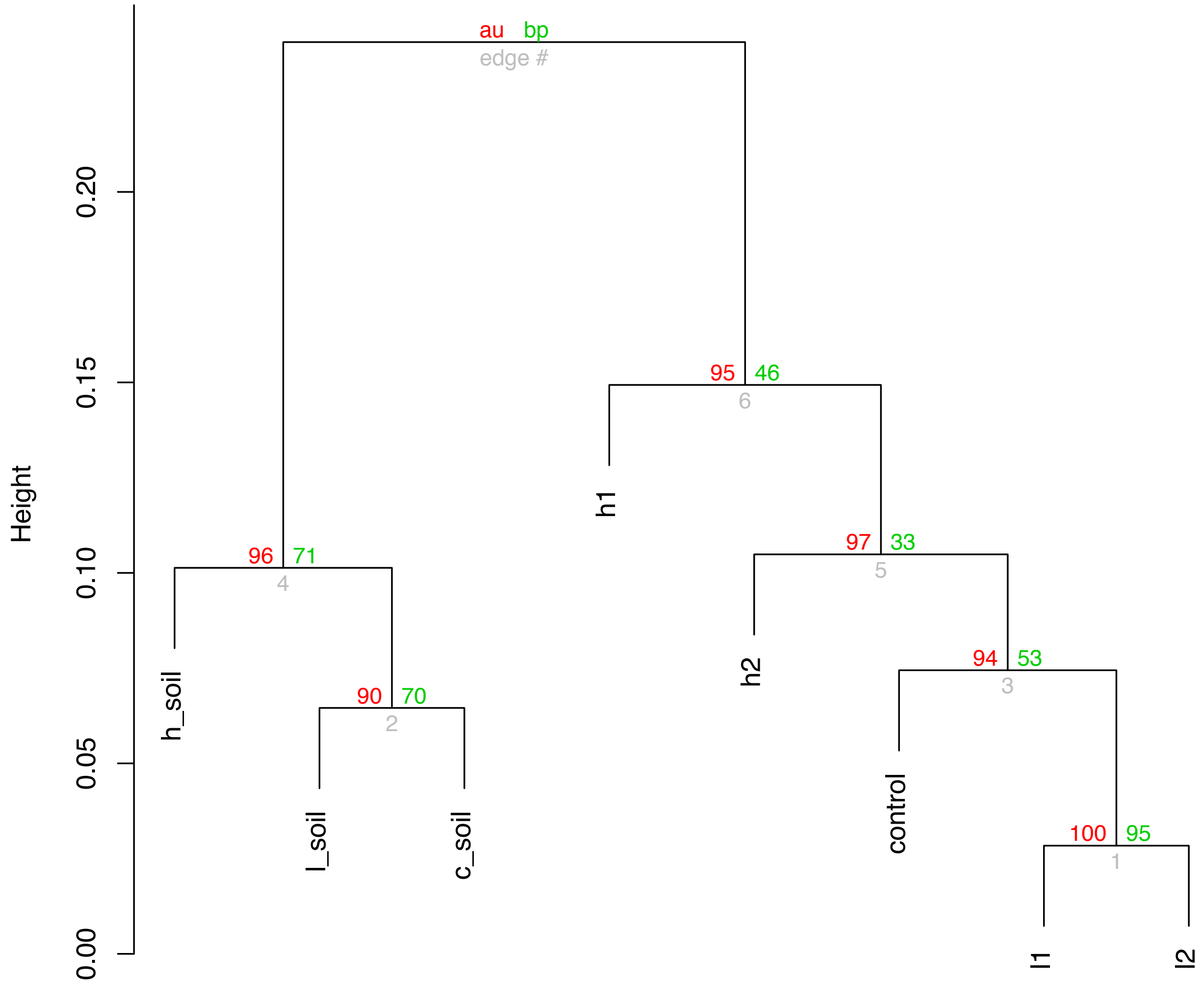


**Supporting Figure 10.** Metabolic capacity for carbon, sulphur, and nitrogen cycles clustered with $P$-values via multiscale bootstrap resampling. AU (approximately unbiased) $P$-values are shown in red, with any value $> 0.95$ (significance level 0.05) being significant. BP (bootstrap proportions) values are shown in green. Hierarchical clustering was completed using "correlation" as distance method and "average" as clustering method.

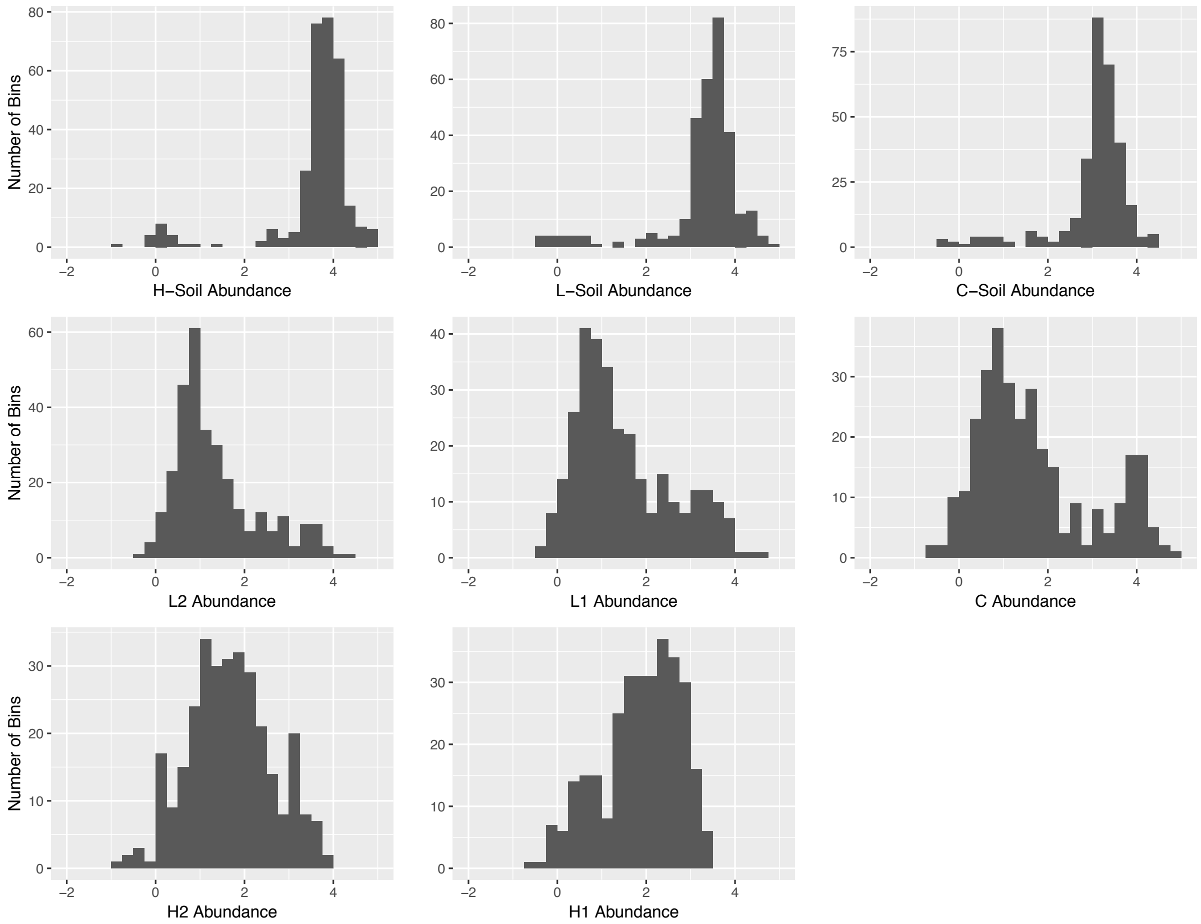


**Supporting Figure 11.** Genome abundance per sample ($-\log_{10}$ scale).

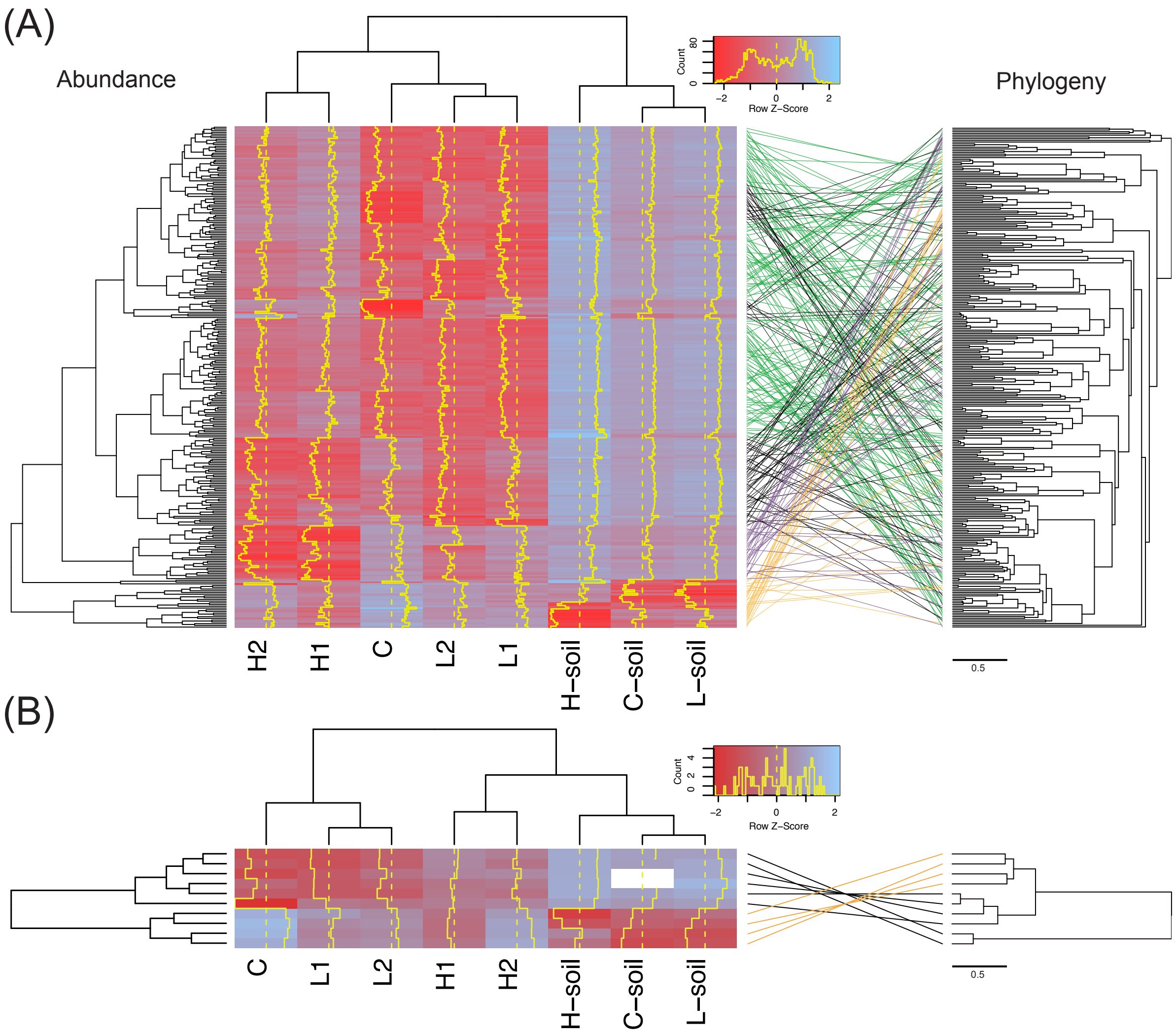


**Supporting Figure 12.** Genome abundance heatmap, and tanglegram separated by (A) bacterial genomes and (B) archaeal genomes. Left: The heatmap displays genome abundance per site normalised by amount of reads in each sample and transformed to a $-\log_{10}$ scale. Dotted yellow lines represent mean abundance values, and yellow traces represent the raw z-scores above (red) and below (blue) the mean. The abundance values are grouped both by sites (top dendrogram) and genomes (left dendrogram). Right: Tanglegram between dendrogram clustered by similar abundances and phylogenetic tree. Highlights in tanglegram: orange lines are genomes abundant in soil, green lines are genomes abundant in low runoff sediment, purple lines are genomes abundant in high runoff sediment, black lines are genomes shared in multiple environments.

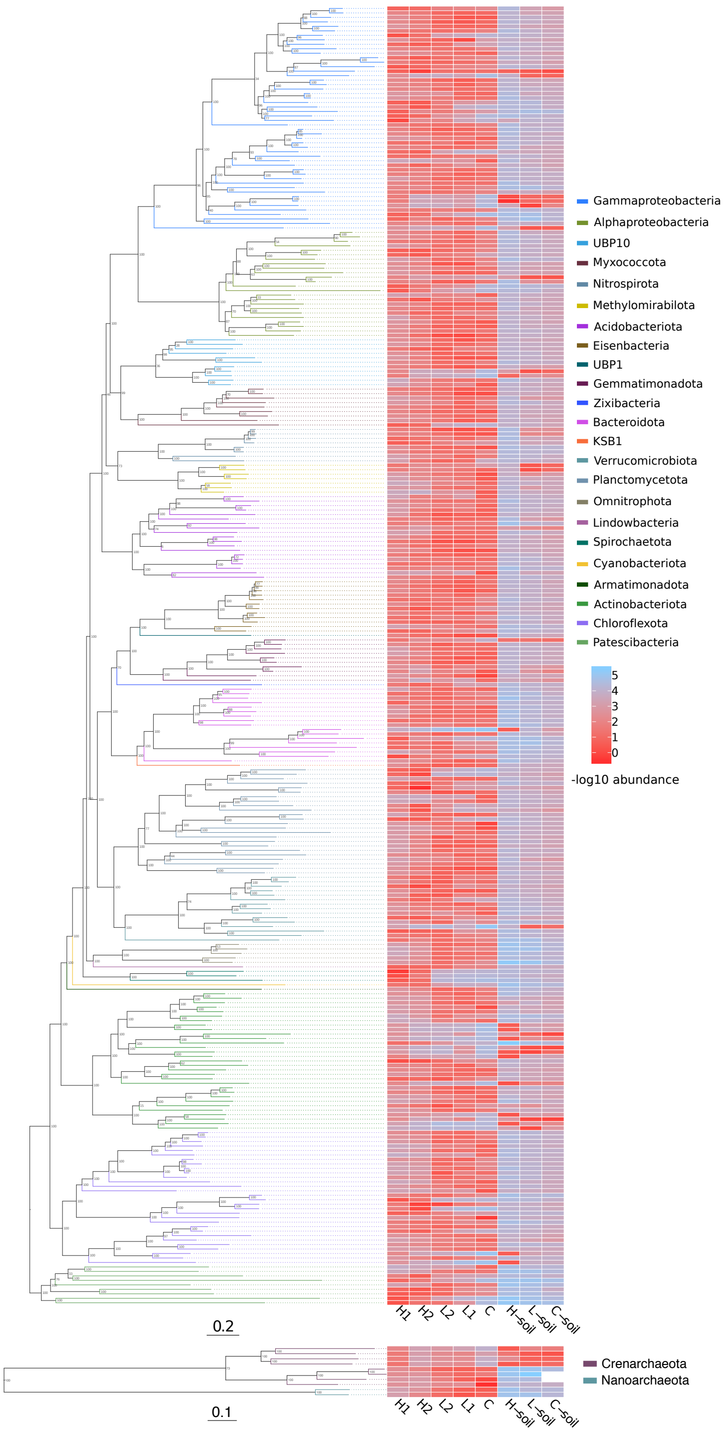


**Supporting Figure 13.** Abundance of genomes are presented on a $-\log_{10}$ scale were more negative values (red) are more abundant than positive values (blue). The abundance values correspond by row with the genomes phylogenetic assignment. Support values for phylogenetic tree are shown at each node.

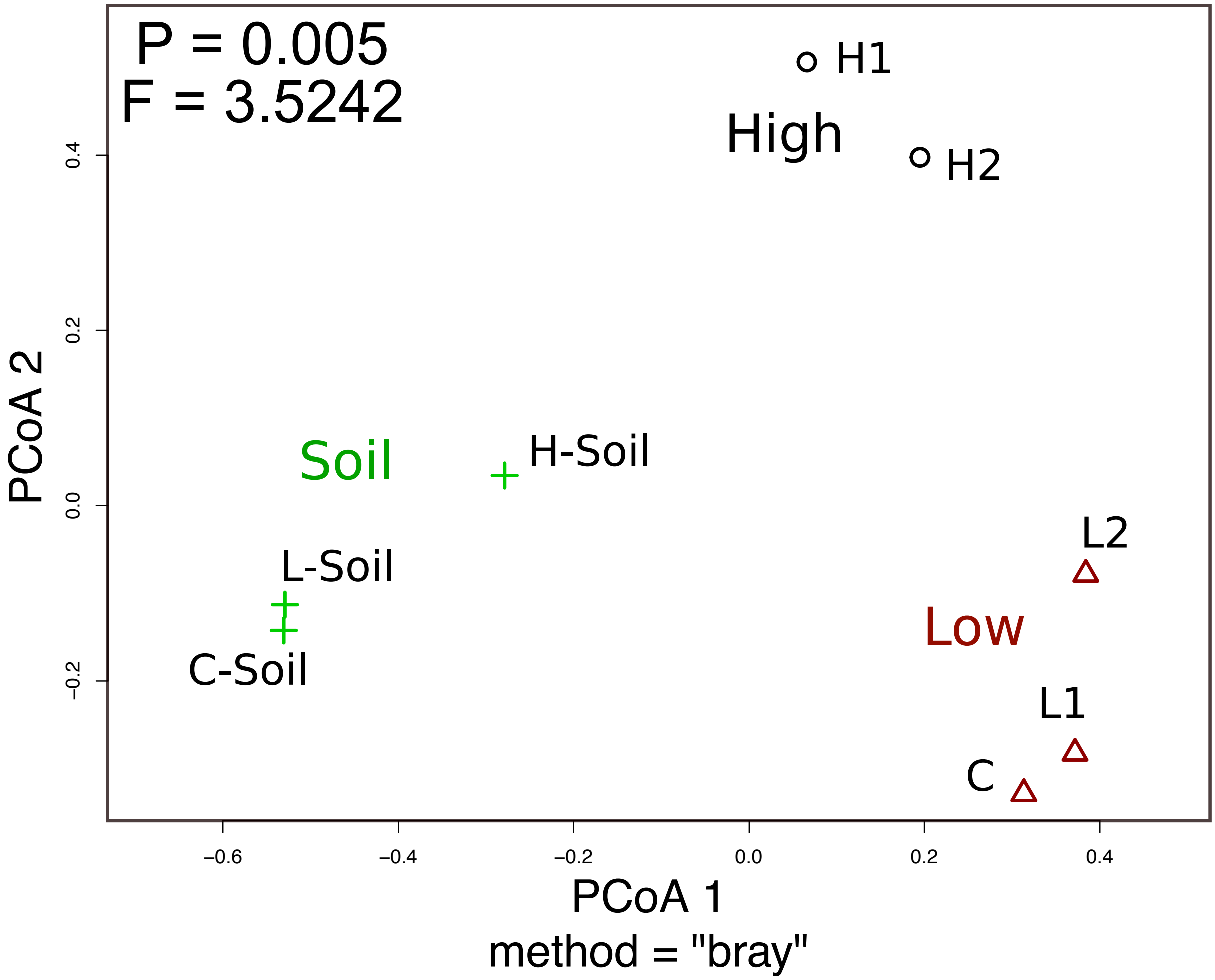


**Supporting Figure 14.** Principal coordinate analysis (PCoA) on reconstructed genome abundances for all sites. Grouping the sites into soil, high, and low runoff was proven to be significant (PERMANOVA test: $F = 3.52$, $P = 0.005$).

### Supporting Tables

**Supporting Table 1.** Coordinates of Lake Hazen sediment and soil sites with temperature and date at time of sampling.

| Sample | Location | Temperature (°C) | Date |
|---|---|---|---|
| **H-Soil** | 81° 84' 840" N; 70° 83' 849" W | -5 | June 3, 2017 |
| **L-Soil** | 81° 80' 332" N; 71° 54' 239" W | 0 | June 7, 2017 |
| **C-Soil** | 81° 79' 382" N; 70° 44' 486" W | -2 | June 3, 2017 |
| **H1** | 81° 84' 150" N; 70° 85' 175" W | not measured | May 24, 2017 |
| **H2** | 81° 82' 493" N; 70° 71' 498" W | not measured | May 29, 2017 |
| **C** | 81° 80' 343" N; 70° 50' 447" W | not measured | May 27, 2017 |
| **L1** | 80° 80' 521" N; 70° 52' 699" W | not measured | June 1, 2017 |
| **L2** | 81° 79' 171" N; 71° 46' 926" W | not measured | June 2, 2017 |

**Supporting Table 2.** Sediment deposition dates and rates for the two deep sites, H2 and L2, is based on $^{210}$Pb constant rate of supply (CRS) dating model. Analysis was completed on 0.5 cm core intervals. Data adapted from previous study [2].

| H2 (Abbe Deepsite) | | | L2 (Blister Deep Site) | | |
|---|---|---|---|---|---|
| **Interval** | **Midpoint CRS date (CRS year)** | **Sedimentation rate (g/cm$^2$/yr)** | **Interval** | **Midpoint CRS date (CRS year)** | **Sedimentation rate (g/cm$^2$/yr)** |
| 0-0.5 | 2017.1 | 0.349 | 0-0.5 | 2016.5 | 0.0724 |
| 0.5-1 | 2016.6 | 0.292 | 0.5-1 | 2014.0 | 0.0701 |
| 1-1.5 | 2015.9 | 0.111 | 1-1.5 | 2011.3 | 0.1163 |
| 1.5-2 | 2014.1 | 0.671 | 1.5-2 | 2009.2 | 0.1759 |
| 2-2.5 | 2012.7 | 0.671 | 2-2.5 | 2007.3 | 0.1232 |
| 2.5-3 | 2012.7 | 0.227 | 2.5-3 | 2004.7 | 0.0909 |
| 3-3.5 | 2012.0 | 5.563 | 3-3.5 | 2001.9 | 0.0967 |
| 3.5-4 | 2011.3 | 5.563 | 3.5-4 | 1998.8 | 0.0854 |
| 4-4.5 | 2011.3 | 5.563 | 4-4.5 | 1994.0 | 0.0513 |
| 4.5-5 | 2011.3 | 5.563 | 4.5-5 | 1987.2 | 0.0375 |
| 5-5.5 | 2011.3 | 5.563 | 5-5.5 | 1976.7 | 0.0220 |
| 5.5-6 | 2011.3 | 5.563 | 5.5-6 | 1966.51 | 0.0515 |
| 6-6.5 | 2011.3 | 5.563 | 6-6.5 | 1957.66 | 0.0339 |
| 6.5-7 | 2011.3 | 5.563 | 6.5-7 | 1949.00 | 0.0537 |
| 7-7.5 | 2011.3 | 5.563 | 7-7.5 | 1940.41 | 0.0451 |
| 7.5-8 | 2011.3 | 5.563 | 7.5-8 | 1931.79 | 0.0668 |
| 8-8.5 | 2011.3 | 5.563 | 8-8.5 | 1925.07 | 0.0553 |
| 8.5-9 | 2011.3 | 0.441 | 8.5-9 | 1918.10 | 0.0553 |
| 9-9.5 | 2010.9 | 0.224 | 9-9.5 | 1911.08 | 0.0553 |
| 9.5-10 | 2009.8 | 0.123 | 9.5-10 | 1904.42 | 0.0553 |

**Supporting Table 3.** Glacial runoff to the Lake Hazen Watershed. Lengths of rivers in km are shown in parentheses. Mass balance modelled runoff for years 2015 and 2016. Sampling dates in 2017 were prior to the summer runoff. Data adapted from previous study [2].

| | **Surface area ($km^2$)** | | | **Runoff volume ($km^3$)** | |
|---|---|---|---|---|---|
| **River** | **Catchment** | **Glacier** | **River** | **2015** | **2016** |
| High Runoff: Abbé (AB) | 390 | 204 | 7.9 (21) | 0.061 | 0.015 |
| Low Runoff: Blister (BR) | n/a | 6 | 2.5 (11) | 0.002 | <0.001 |
| Gilman (GL) | 992 | 708 | 5.1 (22) | 0.192 | 0.08 |
| Henrietta Nesmith (HN) | 1274 | 1041 | 9.6 (4.6) | 0.291 | 0.075 |
| Snowgoose (SG) | 222 | 87 | 5.6 (17) | 0.026 | 0.006 |
| Turnabout (TN) | 678 | 259 | 13.4 (42) | 0.082 | 0.024 |
| Very (VR) | 1035 | 269 | 32.9 (39) | 0.165 | 0.08 |
| Watershed total | 7516 | 3078 | 91.2 | 0.979 | 0.291 |

**Supporting Table 4.** DNA extraction masses. DNA was extracted in triplicate for each sample and then combined prior to sequencing.

| Lake Hazen | Sample Location | Tube ID | Wet Weight (g) | | | | PCR with *glnA* |
|---|---|---|---|---|---|---|---|
| | | | 1 | 2 | 3 | Total (grams) | |
| Sediment | H2: Deephole | S1 | 0.416 | 0.455 | 0.499 | 1.370 | yes (diluted) |
| | L2: Blister Deep | S2 | 0.497 | 0.455 | 0.389 | 1.341 | yes |
| | H1: Abbe | S3 | 0.469 | 0.514 | 0.552 | 1.535 | yes |
| | L1: Blister Shallow | S4 | 0.429 | 0.402 | 0.361 | 1.192 | yes |
| | C: Ruggles | S5 | 0.518 | 0.447 | 0.331 | 1.296 | yes |
| Soil | L-Soil: Blister Soil | BS3 | 0.527 | 0.535 | 0.561 | 1.623 | yes (diluted) |
| | H-Soil: Abbe Soil | AS3 | 0.443 | 0.537 | 0.417 | 1.397 | yes |
| | C-Soil: Ruggles Soil | RS3 | 0.447 | 0.500 | 0.447 | 1.394 | yes |

**Supporting Table 5.** DNA fluorescence assay quantification for each sample. Note: H2 sequencing required using full extraction volume of 65 $\mu$l to reach an appropriate concentration.

| Sample | Volume ($\mu$l) | Concentration ($ng/\mu$l) | Total DNA ($ng$) | NanoDrop (concentration $ng/\mu$l) |
|---|---|---|---|---|
| H2 | 65 | 0.08 | 3.36 | 4.6 |
| L2 | 42 | 2.55 | 107.1 | 7.6 |
| H1 | 42 | 4.33 | 181.86 | 10.2 |
| L2 | 42 | 17.98 | 755.16 | 32.8 |
| C | 42 | 21.2 | 890.4 | 36.4 |
| H-Soil | 42 | 38.14 | 1601.88 | 65.2 |
| L-Soil | 47 | 69.88 | 3284.36 | 149.0 |
| C-Soil | 47 | 74.38 | 3495.86 | 110.0 |

**Supporting Table 6.** Contigs were assembled with Megahit and used in Anvio database. Only contigs that were greater than 2500 bp in length were used to form MAGs, however, contigs with a length of 1000 bp or more were used to form the initial database.

| | Contigs | Total Length (bp) | Min Contig Length (bp) | Max Contig Length (bp) | Avg Contig Length (bp) | N50 (bp) | # of genes prodigal |
|---|---|---|---|---|---|---|---|
| **Megahit** | 12026467 | 8477069127 | 200 | 792468 | 705 | 756 | NA |
| **Anvio** | 1455655 | 3414974759 | 1000 | 792468 | NA | 285024 | 4254625 |

**Supporting Table 7.** Number of bins recovered at each assembly step prior to and after manual refinement with Anvio.

| | **Anvio-Output (CONCOCT binning)** | **CheckM** | **Anvio-Output (manually refined)** | **Final CheckM** |
|---|---|---|---|---|
| Total Bins | 850 | 850 | 877 | 877 |
| Completion, Redundancy | | | | |
| C: >90, R: <10 | 52 | 74 | 52 | 72 |
| C: >70, R: <10 | 180 | 199 | 178 | 198 |
| C: >50, R: <10 | 321 | 324 | 309 | 300 |
| C: <50, R: any | 529 | 526 | 568 | 577 |
| Contamination | | | | |
| <1 | 233 | 261 | 255 | 286 |
| <5 | 544 | 632 | 590 | 676 |
| >10 | 8 | 51 | 7 | 27 |
| Good bins but contaminated | | | | |
| C: >50, R: >10 | 2 | 27 | 0 | 2 |
| C: >70, R: >10 | 1 | 6 | 0 | 0 |
| C: >90, R: >10 | 0 | 2 | 0 | 0 |

**Supporting Table 8.** *P*-values for marker gene and pathway distribution between sites. A Fisher's exact test was used in the place of $\chi^2$ test for small count numbers. *P*-values of less than 0.05 are considered significant (labelled with an asterisks).

| Functional Marker | High vs Low | High vs Control | High vs Soil | Low vs Control | Low vs Soil | Control vs Soil |
|---|---|---|---|---|---|---|
| Carbon Cycle | 0.03298 * | 0.0004998 * | 0.0004998 * | 0.07746 | 0.1384 | 0.001499 * |
| Nitrogen Cycle | 0.07596 | 0.001999 * | 0.0004998 * | 0.7006 | 0.03448 * | 0.1859 |
| sulphur Cycle | 0.001999 * | 0.002999 * | 0.0004998 * | 0.4913 | 0.02999 * | 0.3823 |
